# The *Multi-scale Copula Index (MCI)*: a new information-theoretic measure of association and its applications in functional cancer genomics

**DOI:** 10.64898/2026.10.10.758174

**Authors:** Timothy Pham, Karina Meneses-Cime, Jiayan (Yoshii) Ma, Bartholomew Tam, Amey Gupta, Michael Reich, Alex Wenzel, David Eby, Thorin Tabor, Helga Thorvaldsdóttir, Antony Castanza, Jill P. Mesirov, William Kim, Pablo Tamayo

**Affiliations:** Moores Cancer Center, UC San Diego, La Jolla, CA, USA; UC San Diego School of Medicine, La Jolla, CA, USA; Eli and Edythe Broad Institute of Harvard and MIT, Cambridge, MA, USA; Yale University School of Medicine, New Haven, CT, USA

## Abstract

Cancer genomic analyses increasingly involve large, complex datasets representing diverse ‘omics’ variables, such as gene expression, mutations, copy number alterations, genetic perturbations (e.g. CRISPR or iRNA), drug sensitivity, and clinical phenotypes, making it crucial to accurately measure associations among these variables. Traditional correlation-based methods, like the Pearson correlation coefficient, often assume linearity, Gaussian distributions, and independence among variables, leading to oversimplified assumptions, less sensitive readouts and less accurate significance estimation. Information-theoretic measures overcome these limitations by being one step removed from the data and assessing dependencies at the level of probability distributions, thus providing a more data-agnostic, robust measure for identifying associations in common analysis tasks such as feature selection, pattern recognition, and predictive modeling. Building on these foundations, we introduce the Multi-scale Copula Index (*MCI*), a novel information-theoretic measure of association. The *MCI* offers significant advantages over existing methods by eliminating the need for density estimation and parameter tuning, thereby ensuring consistent estimation with reduced computational complexity. Furthermore, it enhances statistical power at lower sample sizes, exhibits strong robustness to outliers, and successfully captures both directionality and important symmetries under input-data transformations, such as reflection or complementation. Due to these strengths, the *MCI* is particularly useful for functional cancer genomic applications involving, e.g., transcriptional and perturbational data, with different statistical properties across a wide range of dataset sizes and complexity. Contextual associations often arise in functional genomics datasets: two genomic variables show little or no correlation in most samples, yet display a strong linkage within the subset that reflects the “active” biological context. These situations are ideal for quantifying associations with the *MCI*. We illustrate the performance of the *MCI* in a variety of benchmarks and cancer genomic examples and applications. We also introduce the *Cumulative Entropy Diagram* (CED), a graphical display that allows the visualization of the *MCI* between a reference vector, such as a phenotype, pathway activation or drug response, and a complete set of genomic features.

## Introduction and Motivation

Functional cancer genomic analysis today routinely involves larger, more complex datasets and the integration of diverse ‘omics data types [1–5]. In this context, researchers need to compare genomic variables with different statistical properties—such as gene expression numerical profiles, CRISPR/Cas9 perturbations, drug sensitivity measurements, and biological or clinical phenotypes—to establish their degrees of association. This analysis typically involves matching numerous genomic variables against selected phenotypes and identifying top hits as candidates for further study or as inputs to predictive models [6,7]. These comparisons often reveal complex relationships between variables, reflecting the contextual nature of cellular networks. Despite their fundamental importance, feature selection and association analysis methodologies frequently rely on rudimentary statistical algorithms that make oversimplistic assumptions about the data’s properties. For instance, the standard Pearson correlation assumes linear relationships between genomic feature vectors, yet biological data often exhibits non-linear associations that cannot be accurately quantified under linearity assumptions. Similarly, other simplistic methods presume Gaussian distributions of genomic readouts and gene independence, resulting in serious overestimation of statistical significance, inflated p-values, and spurious conclusions [8]. Even traditional power calculations for determining sample sizes require reassessment and improvement in the context of large genomic datasets [9]. Other more modern measures, such as the distance correlation (dCor [9]) and the Hilbert-Schmidt Independence Criterion (HSIC[10]), are better at detecting non-linear associations but rely on computing pairwise distances or kernel similarities between samples resulting in a time and memory complexity of O(N^2^) for a dataset with N samples. In addition those methods provide no directionality of association. Other sophisticated approaches based on Information-Theoretic measures, provide more powerful and sensitive methods but they pose challenges that have limited their practical use in genomics analysis. Mutual information (MI), and other variants of it such as MIC[11], requires knowing the underlying probability distributions of the variables, the simplest way to estimate being to divide the data into discrete bins to form a histogram. However, the choice of bin size and number can drastically alter the results. To avoid binning, researchers use methods like Kernel Density Estimation (KDE [12]) or k-Nearest Neighbors (k-NN [13]) estimators, however, these methods are vastly more computationally intensive than simple histograms. Those information theoretic approaches, including HSIC, and other kernel-based methods, also require the tuning of parameters, such as the kernel bandwidth adding complexity to the estimation method. Given these limitations, continued development and application of more sensitive, data-agnostic association measures are essential to better model the diverse types of genomic data with varying statistical properties in increasingly large datasets.

### A genomic association example

We introduce here a new association measure to address those challenges based on the Cumulative Copula Entropy [14,15] that we refer to here as the Multi-scale Copula Index (MCI). We first motivate its usefulness and provide a concrete example and then describe THE MCI in detail. The example include twelve instances of statistical dependency between mRNA and CRISPR/Cas9 standardized profiles for the same oncogene across 1,112 cancer cell lines from the Dependency Map^1^ (depmap.org). This is shown in Fig. 1 where scatter plots show the knockdown sensitivity values (CRISPR/Cas9) vs. gene expression (mRNA) for 12 oncogenes in the 1,112 cancer cell lines that are common to both datasets.

**Figure 1.**
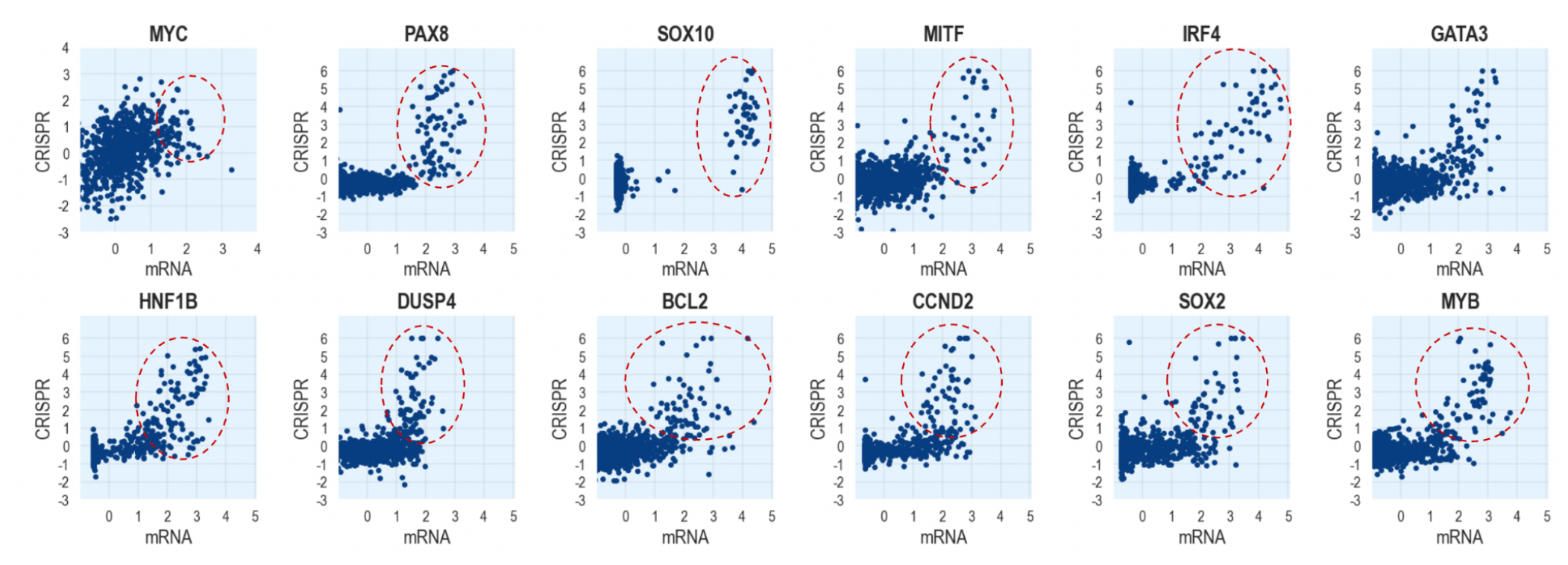
Examples of relationships between CRISPR and mRNA profiles for 12 selected oncogenes in 1,112 cancer cell lines.

The relationships between a gene’s CRISPR knockdown sensitivity and its corresponding gene expression profile are important in functional genomics analysis, as they can indicate “oncogene addiction,” where cancer cells become highly dependent on the ongoing activity of a specific oncogene, sometimes mutated, but not always, for their growth and survival [16,17]. In Hahn et al. (2021) [6], we advocated for the study of cancer genes exhibiting this behavior as a means to expand the number and diversity of cancer therapeutic targets. In Fig. 1, it is evident that the relationships between the CRISPR and mRNA profiles of these oncogenes are complex, non-linear, and display a range of diverse statistical behaviors, such as multimodality and long tails. This is particularly conspicuous in the upper right hand side behavior of the plots, where data points exhibit an upward tail (red dashed-line regions). It appears that there is a well-defined context in which above-average gene expression values correspond to higher CRISPR sensitivity profiles, presumably reflecting one or more cellular states where the oncogene is active or induced, making cancer cells more dependent on it. Outside of that specific context, such as in the bulk of the distributions, the association is less pronounced. We now analyze in more detail the first two oncogenes, MYC and PAX8 as shown in Fig. 2.

**Figure 2.**
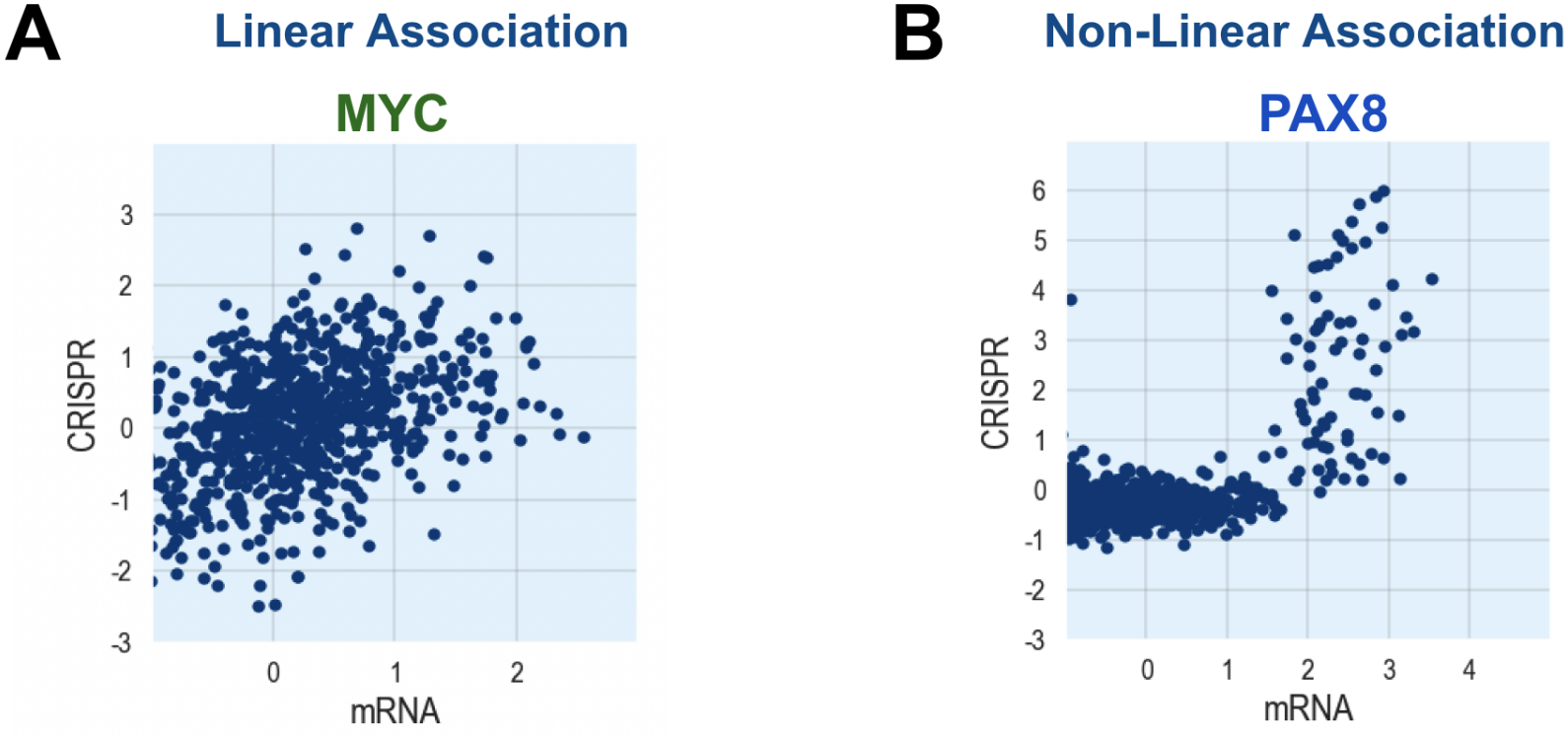
Relationships between CRISPR knockdown and mRNA profiles for the MYC (A) and PAX8 (B) oncogenes.

As can be seen in Fig. 2A, the CRISPR vs. mRNA association for MYC is more linear-like and the CRISPR knockdown appears to involve a broad effect on the viability of many cell lines where higher sensitivity roughly correlates with higher expression and vice versa. In this case simple measures of association, such as the Spearman coefficient, are able to detect and effectively quantify this relationship. In Fig. 3A we match the entire genome-wide dataset of CRISPR profiles against the mRNA sensitivity profile of MYC and we can see that the MYC CRISPR profile is for both measures the top match to the MYC mRNA target, out of a total of 15,778 genes in 1,112 samples, and the results from using both measures are practically the same overall. As we will see below (eq. 30), the *MCI* measure is normalized to approximate the Spearman coefficient for Gaussian distributed vectors with linear association and this is the reason both measures produce essentially the same results in this example. For situations like this, where most of the relationship is linear, the *MCI* measure is roughly equivalent to the Spearman correlation.

**Figure 3.**
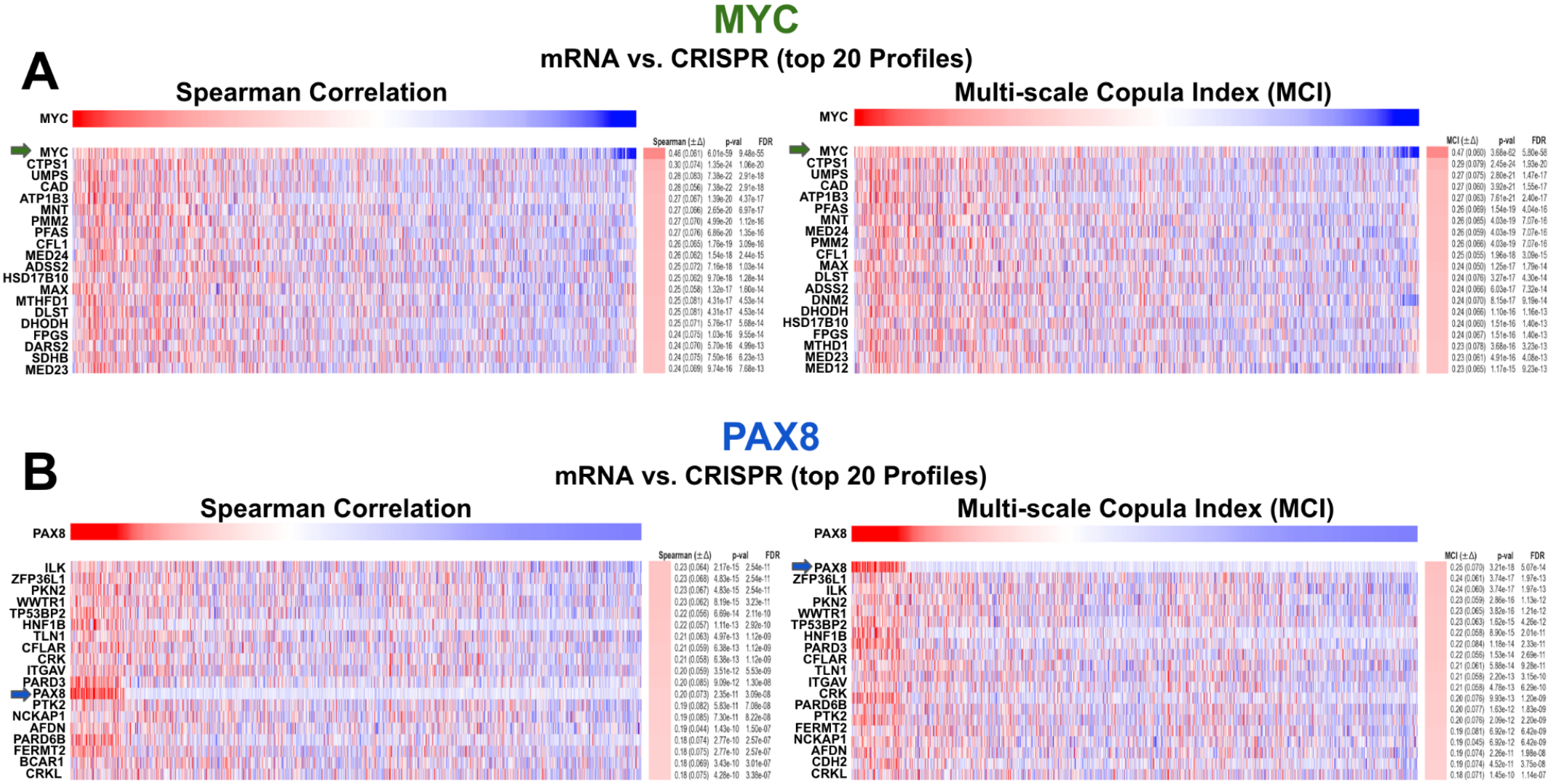
Top 20 CRISPR profiles matching A) MYC, and B) PAX8 mRNA profiles using the Spearman correlation and the MCI.

In contrast, in the case of PAX8 (Fig. 2B), the CRISPR vs. mRNA association is very non-linear and the sensitivity of many cell lines appear to be low and mostly independent of the level of expression until a threshold, of about 1.5 standard deviation above the mean, is reached and then the CRISPR knockdown sensitivity increases significantly. This very non-linear behavior of PAX8 is not fully quantified by simple association measures such as Spearman correlation. In Fig. 3B, where we match the CRISPR profiles against the mRNA profile of PAX8, we can see that the PAX8 CRISPR is number 12 down the list for the Spearman correlation, however it is the top match according to the *MCI* measure. More quantitatively, we can see in Fig. 4 that for MYC both measures and their statistical significant estimates (p-values and FDRs estimates) are equivalent, however for PAX8 the *MCI* estimate is 25% higher, 0.25 vs. 0.20, and also attains much more significant statistical significance in terms of p-values and FDRs. The *MCI* measure is not only more sensitive, but the lower p-values and FDRs indicate that it also has higher specificity to detect non-linear associations^2^.

**Figure 4.**
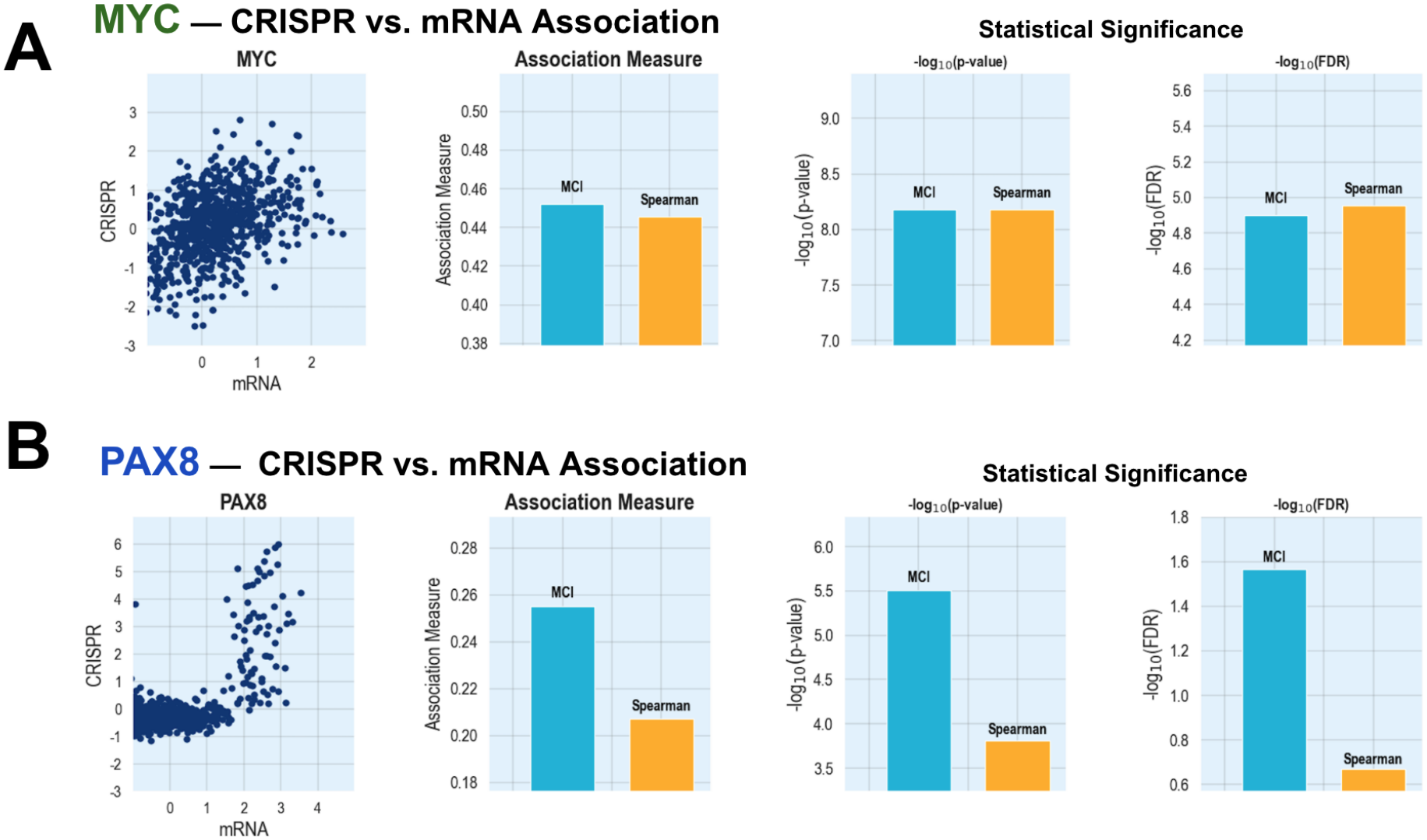
A) MYC and B) PAX8 mRNA vs. CRISPR association and statistical significance (p-values and FDRs).

This example shows how the *MCI* measure of association can detect non-linear relationships and offer better sensitivity and specificity not only in assessing isolated relationships such as those shown in Fig. 1 but also in selecting the best matching features from a large universe of candidates as shown in Fig. 3.

It is important to mention that other information-theoretic measures can also effectively achieve this higher sensitivity and specificity when compared with simpler and more traditional measures. For example, we have used the Information Coefficient (*IC*) [18,19], a rescaled version of the mutual information, and the corresponding Conditional Information Coefficient (*CIC*) [20,21], for over a decade to provide sensitive feature selection methods to find associations between genomic variables [20,22–25]. While the *IC* has proven effective, its use in specific datasets presents several challenges. Accurately approximating the *IC* with kernel density estimation (KDE) requires selecting appropriate bandwidths for the variable distributions—an especially difficult task for small or medium-sized datasets. Moreover, the bandwidth must be adjusted based on the correlation between variables to prevent overestimation at low correlation values and underestimation at high ones. Additionally, both bandwidth selection and KDE are computationally intensive, making the *IC* calculation over 30 times slower than standard correlation measures such as the Pearson and Spearman correlation. This significantly limits its scalability, particularly for large datasets or when using outer-loop iterative procedures such as all variables vs. all variables association or permutation-based testing, etc.

Addressing these specific challenges while preserving the advantages of an information-theoretic measure for detecting non-linear associations was the primary motivation behind the development of the Multi-scale Copula Index (*MCI*). This measure can be understood as part of the broader category of information-theoretic measures of dependence, similar to mutual information, but it offers certain computational and mathematical advantages that overcome the limitations described above. It is based on copula statistics [26–28], which are mathematical tools that enable the modeling and analysis of the dependence structure between random variables, separate from their individual distributions. Copulas are particularly valuable because they capture complex relationships between variables that extend beyond simple linear correlation, including tail dependencies and asymmetric associations. Sklar’s theorem serves as the theoretical foundation for copula-based methods in statistics, finance, risk modeling, and many other fields where a deeper understanding of dependencies between variables is crucial [26–28]. This theorem states that any multivariate distribution can be separated into two parts: the individual behavior of each variable, represented by the individual marginal distributions, and the dependency structure between variables, represented by the copula function. This separation is powerful because it allows us to model dependencies independently of their marginal distributions and to study them without being influenced by the specific characteristics or idiosyncrasies of individual variables. In data analysis, copulas are essential in any scenario where understanding how variables associate with one another is more important than examining their individual behaviors [26–28].

Now we will review the relevant building blocks necessary to understand and define the *MCI* such as mutual information, cumulative entropy, copulas and cumulative copula entropy. If the reader is already familiar with mutual information, copulas and cumulative entropies, he or she may want to skip the next 3 sections and go directly to the *Defining the Multi-scale Copula Index (MCI)* section below.

### Entropy, Mutual Information and the Information Coefficient

In what follows we use the standard notational convention that uses capital letters (*X, Y, U, V*, etc.) to represent random variables themselves, and lowercase letters (*x, y, u, v*, etc.) to represent specific values that these random variables can take. In information theory entropy, *H(X)*, quantifies the average amount of “information” and is a measure of a random variable’s uncertainty [29–31]. For a continuous random variable *X*, it is,

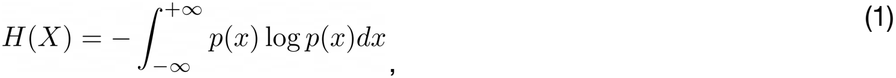

where *p(x)* is the probability density of *X*. The maximum entropy occurs when all values or outcomes are equally likely, and the entropy is zero when there is complete certainty about the value or outcome of the variable. Entropy captures a crucial concept in applications such as data compression and communication, as it quantifies information and establishes limits on the efficiency of encoding it. It is also useful in data analysis as it quantifies the amount of information in observed or measured empirical variables such as a genomic readout or a phenotype. The concept can be extended to two continuous variables, *X* and *Y*, where their joint entropy is a function of their joint probability density, *p*(*x, y*),

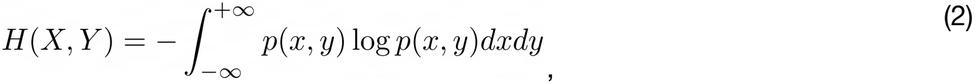

Mutual information is a fundamental concept in information theory that quantifies the amount of information shared between random variables [29–31]. It measures how much knowledge of one variable reduces the uncertainty about the other. This is relevant in applications where we investigate the degree of association of one genomic variable, or phenotype, with another. The mutual information is defined as the difference between the variables’ individual and joint entropies,

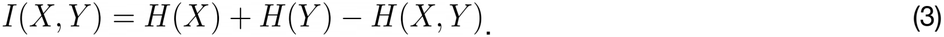

It is also the average log ratio between the variables’ joint and marginal probability densities,

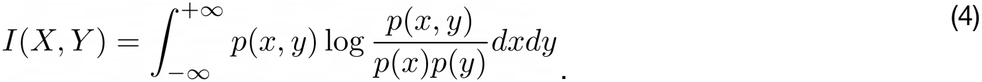

When the variables are independent, the joint distribution is the product of the marginal distributions, so the mutual information is zero. Other important properties of the mutual information include the fact that it is always non-negative and that it is symmetric under exchange of the arguments (*I*(*x, y*) = *I*(*y, x*)).

Mutual information captures both linear and nonlinear dependencies, making it a versatile and powerful measure of association in data analysis, statistical modeling, and machine learning. One of its limitations is that it has a finite lower bound of zero for independent variables with no correlation, while the upper bound is determined by the variable’s entropy, which varies for each variable. One way to address this issue is to rescale mutual information so that it falls within the range [0, 1], similar to the correlation coefficient. This can be achieved using the relationship between mutual information and the correlation coefficient for Gaussian-distributed variables, resulting in Linfoot’s Information Coefficient (*IC*) [18,19][18,19][18,19],

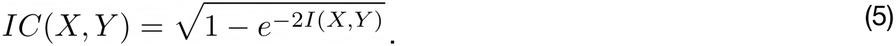

The *IC* can also be made signed and sensitive to the directionality of association, by giving it the sign of the correlation coefficient *ϱ* (*x, y*) between the variables,

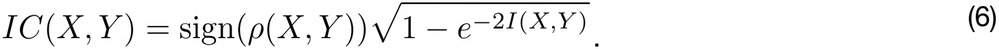

This signed and rescaled version of the mutual information is defined in the interval [-1, 1] like the standard correlation coefficient, and is very useful in practical applications. We have made an efficient implementation of the *IC* (eq. 6) using kernel density estimation^3^ and used it in many cancer genomics studies to measure the association between phenotypes and genomic features, e.g., in the study of gastrointestinal stromal tumors [32], head and neck cancers [33,34], myelodysplastic syndromes [35], prostate cancer [36,37], medulloblastoma [38], and melanoma [39,40]. We have also implemented a conditional version of the *IC*, the *Conditional Information Coefficient* (*CIC*) to find groups of mutually exclusive genomic alterations that explain a response phenotype better than any individual alteration alone and rarely co-occur in the same tumor [20,21].

### Cumulative entropy

The cumulative entropy, *CE*(*x*), is defined analogously to the differential entropy [29,31] but using the cumulative distribution function *P*(*X ⩽ x*) instead of the probability density [41,42],

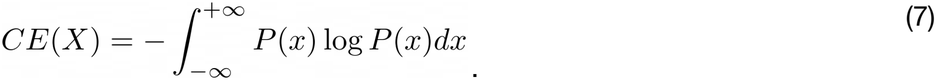

For two variables *x* and *y*, the joint cumulative entropy, *CE(x, y)*, is,

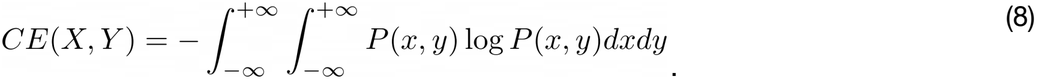

Cumulative entropy offers several advantages. For instance, it is always non-negative, unlike standard differential entropy, which can be negative, for example, in the case of a uniform distribution with very narrow support or a Gaussian distribution with a small standard deviation (2πeσ^2^ < 1). Cumulative entropy is consistently non-negative and arguably provides a better representation of uncertainty or information content than differential entropy. Additionally, cumulative entropy offers a definition that is valid and consistent for both continuous and discrete random variables [42], while standard Shannon entropy requires separate definitions for these types. Moreover, discrete entropy does not naturally converge to differential entropy in the continuous limit. Cumulative entropy can also be defined in cases where the probability density does not exist, making it more general [43] and easier to estimate from empirical data using data-fitted empirical distribution functions [42].

### Copulas and Copula Entropy

Sklar’s theorem is a fundamental result in multivariate statistics that establishes the connection between multivariate distributions and their underlying statistical dependencies. The theorem states that any multivariate joint cumulative distribution function (CDF) *F(X*_1_,*X*_2_, …, *X*_*n*_), can be defined as a function of its marginal distributions *F*_1_*(X*_1_*), F*_2_*(X*_2_),…, *F*_*n*_*(X*_*n*_), using a *copula*^4^ [26–28] function *C*(…),

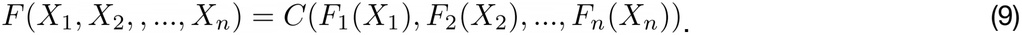

This also implies that there is a copula density *c*, the derivative of the copula function [15],

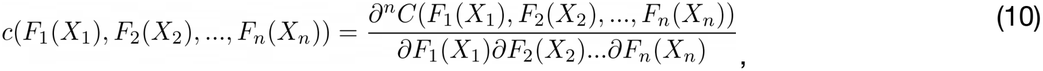

that allows decomposition of the multivariate density *f(X*_1_, *X*_2_, …, *X*_*n*_) as a product of the copula density times the marginal densities,

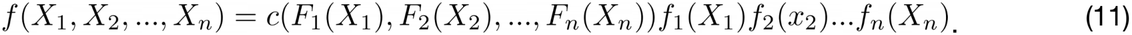

Moreover, if all of the marginal distributions are continuous, then this copula function is unique. If we make a change of variables *U*_*1*_ *= F*_*1*_(*X*_*1*_), *U*_*2*_ *= F*_*2*_(*X*_*2*_),…, *U*_*n*_*=F*_*n*_(*X*_*n*_) we can see that the copula *C*(*U*_*1*_, *U*_*2*_,…, *U*_*n*_) can be defined over new variables (*U*_*1*_,…, *U*_*n*_) with uniform distributions on the product of intervals [0,1]^n^. Ma and Sun showed that the entropy of a bivariate copula density between two variables *X* and *Y*, is the negative of the mutual information [44,45]. As we saw above, the mutual information between random variables *X* and *Y* is defined as:

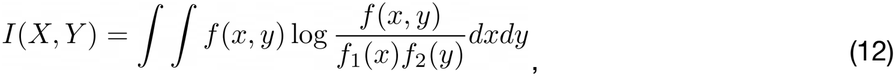

where *f*(*x,y*) is the joint density function and *f*_*1*_(*x*) and *f*_*2*_(*y*) are the marginal densities. By Sklar’s theorem, we can express the joint density as,

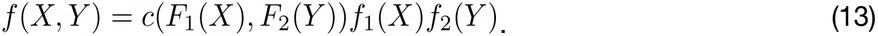

Where *c*(*F*_*1*_(*x*), *F*_*2*_(*y*)) is the copula density and *F*_*1*_(*x*) and *F*_*2*_(*y*) are the marginal CDFs. If we substitute this into the mutual information (eq. 12),

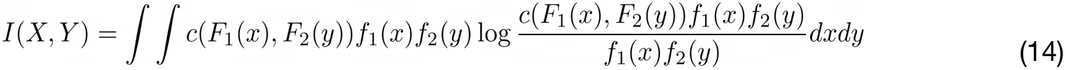

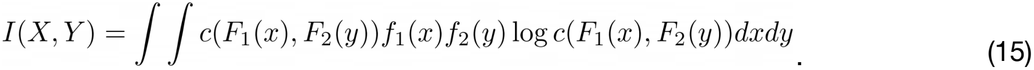

Making the variable transformation *U = F*_*1*_(*X*) and *V = F*_*2*_(*Y*), which have uniform distributions on [0,1], and the fact that the probability densities are the derivatives of the marginal CDFs: *f*_*1*_(*x*) *dx = dF*_*1*_(*x*) *= u* and *f*_*2*_(*y*) *dy = dF*_*2*_(*y*) *= v*, we derive mutual information in terms of the copula density,

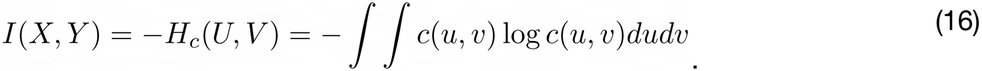

Therefore, the entropy of the copula density is the negative of the mutual information between *X* and *Y*,

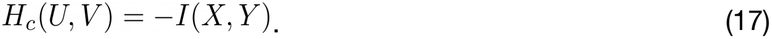

The entropy of the density copula, *H*_*c*_(*u, v*), can be viewed as an embodiment of the shared information between two variables, specifically representing pure “association information” that is completely detached from the information contained in the marginal distributions [44,45]. In this way, the joint entropy can be understood as the sum of the marginal entropies plus the copula entropy,

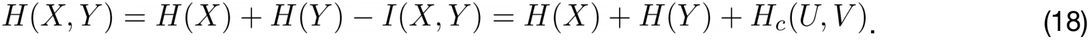

The key advantage of copula entropy is that it combines the powerful dependency detection of an information-theoretic measure with the marginal-free property of copulas, making it both theoretically elegant and practically useful for complex data analysis. Interestingly, despite the introduction of copula theory in 1959 and mutual information in 1948, it seemingly took almost 50 years for researchers to recognize the striking connection between the two mathematical concepts [45].

### The Cumulative Copula Entropy

The remarkable connection between the copula density entropy and the mutual information has motivated the search for other information-theoretic measures of dependency based on copulas. The *Cumulative Copula Entropy* (*CCE*) was recently introduced by Nair and Sunoj [14] and Arshad et al. [15], to tackle several limitations of estimating dependencies via the copula density (eq. 10). The *CCE* is defined using the copula function *C*(*U, V*) (eq. 9) as,

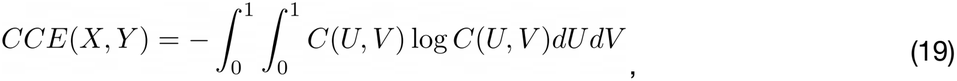

where as before, *U = F*_*x*_(*x*), and *V = F*_*y*_(*y*). The cumulative copula can be defined from the copula density in two dimensions *c*(*u, v*) by performing integration,

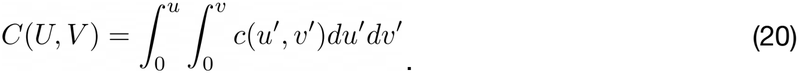

The *CCE* is a function of the cumulative copula *C*(*U, V*), which is in turn based on the cumulative distributions (*U, V*). Therefore, it serves as the copula-based counterpart to the cumulative entropy (eq. 8) [41,46]. As such, it inherits some advantages of using cumulative distribution functions (CDFs) rather than probability density functions (PDFs), including the fact that empirical CDFs converge uniformly to the true CDF according to the Glivenko-Cantelli theorem [47]. Consequently, the *CCE* offers stronger statistical guarantees than density estimators. The integrand in eq. 19, which we refer to as the “*entropula*,” can be thought of as a local function *ℰ* (*u, v*) that measures the amount of “local information” in the cumulative copula,

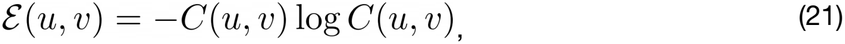

making the *CCE* the expected value of the *entropula*,

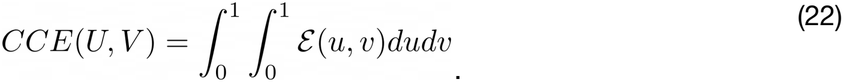

As a measure of dependence, the *CCE* also offers practical advantages [14,15]. For example, it demonstrates improved robustness to noise because it is based on distribution functions, making it more stable when estimated with relatively small samples. Additionally, working with cumulative distributions eliminates the need for density or bandwidth estimation, thereby reducing the associated computational burden and enabling significantly faster implementations. The *CCE* represents a significant advancement in the search for better multivariate measures of dependence. However, despite its many advantages, it can be further improved in various ways to enhance its theoretical properties and practical implementation characteristics, making it better suited for numerous application domains, including functional cancer genomics. For instance, it can be made multi-scale to capture statistical dependencies at multiple levels of resolution. Additionally, it can be defined to possess extra symmetries with respect to order-reversing transformations, and intrinsic directionality, allowing it to detect both positive and negative associations. Finally, it can be rescaled to be bounded within the interval [™1,1] and be numerically rescaled to approximate the Spearman coefficient for Gaussian variables with linear association. In the following sections, we will describe the *Multi-scale Copula Index* (*MCI*) and its properties.

### Defining the Multi-scale Copula Index (*MCI*)

In what follows we will consider the association between two random variables *x* and *y* and will consider bivariate copulas but the definitions can easily be generalized to *n* dimensions. The *CCE* is symmetric with respect to the reversal of the input variables, namely,

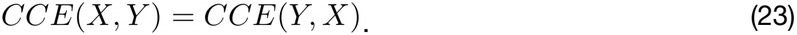

However, it lacks symmetry with respect to order-reversing transformations, such as reflection (*x → -x*) or complementation (*x → max*(*x*) *-x*), of one of the input variables. This specific symmetry represents an important and desirable property in many applications where, if we reverse the order of one variable through reflection or complementation, we would expect to obtain the same degree of association as before but with a negative sign. This behavior also requires that the measure is able to detect negative association and therefore possess directionality, so that the reverse ordering transformation produces the same measure but with opposite sign. This is a common scenario in cancer genomics where researchers associate, for example, a phenotype *x*, such as response to a drug, with a genomic variable *y*, such as the status of a specific gene mutation or the mRNA profile of a gene. If we reflect or complement the phenotype, e.g. replace the wild type with mutant status, it is desirable to obtain the same measure of association but with a negative sign. In summary, we require that our new measure *MCI*(*x, y*), displays the following symmetries,

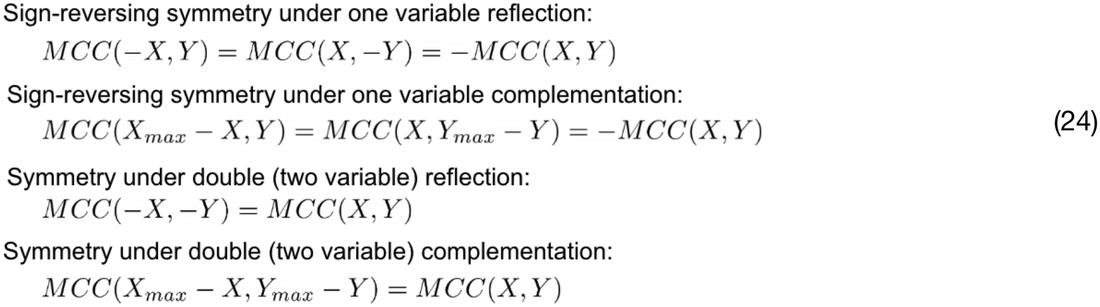

We can impose these symmetries, and at the same time introduce directionality to the measure, by combining the different versions of the cumulative copula functions,

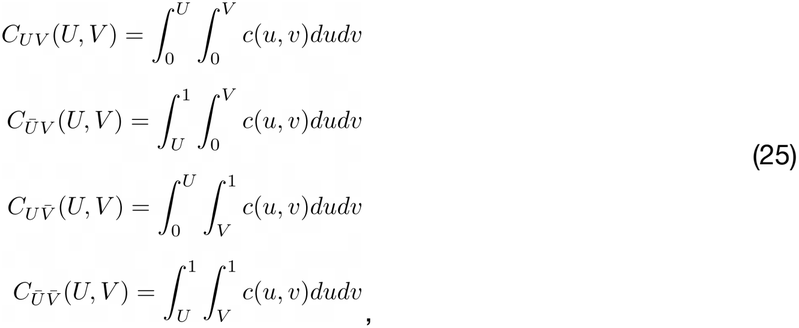

where the bar over the variables indicates the integration is in the backward direction. The first copula is the standard cumulative copula and the last is the survival or reliability copula, and the other two are “hybrid” copulas. The corresponding local entropies, i.e. the *entropulas*, are then,

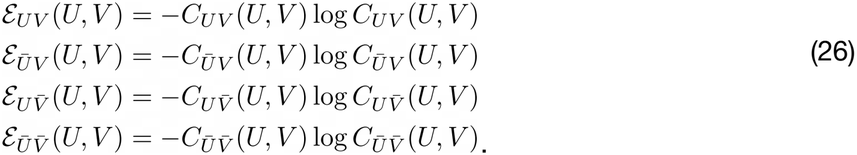

These functions represent the “local” information in each of the different copulas. The corresponding *CCE* entropies for these copulas are then,

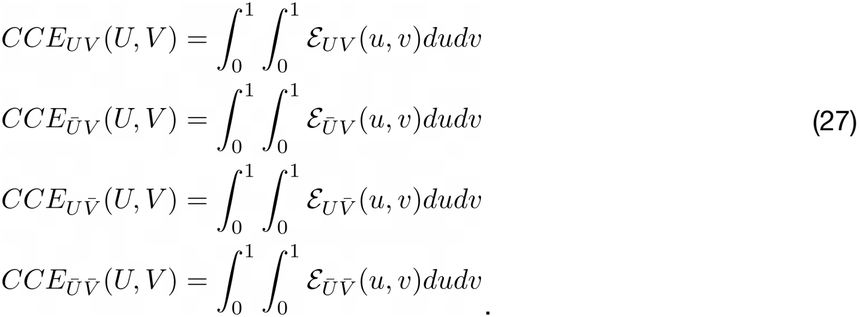

We define the *Symmetrized Copula Cumulative entropy* (*SCCE*) as the difference between the *CCE* entropies of the standard cumulative and survival copulas, minus the hybrid ones,

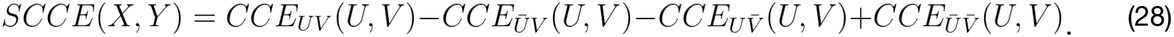

This combination of *CCE* entropies provides directionality because the measure can detect negative correlations when the hybrid copulas’ cumulative entropies are larger than those for the direct copulas. This expression also produces the desired symmetries corresponding (eq. 24) because the effect of reflection or complementation is to reverse the ordering of the data along one of the cumulative copulas’ axes. After an order-reversing transformation in one of the inputs, the value of the *CCE* will have the same magnitude but opposite sign (sign-reversing symmetry) as before the transformation as we show in detail in the appendix. The pairing of the cumulative and survival copula entropies in the *SCCE* is similar to the pairing of the cumulative and survival entropies in Klein and Doll’s cumulative paired entropy [48]. An additional improvement comes from estimating the *SCCE* hierarchically at different scales, for example, by estimating it using computational lattices of different sizes (see details in the next section). This allows us to detect association at a baseline level but also at more coarse-grained scales. The motivation for this comes from our previous experience designing multi-grid methods in Statistical Physics [49,50], and also is one of the properties of the *Maximal Information Coefficient* (*MIC*) [11]. Taking all this into account, it is desirable to define a multiscale *SCCE* and then define a *Multi-scale Symmetrized Cumulative Copula Entropy* (*MSCCE*) that takes the maximum association as measured by the *SCCE*, positive or negative, across different scales (*g*_*min*_,*…,g*_*max*_),

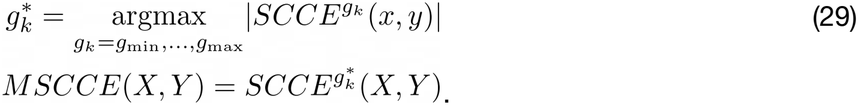

Finally, the *MSCCE* is rescaled to approximate the Spearman correlation for Gaussian distributed inputs. A natural way to accomplish this would be to use the known bounds on the cumulative copula [51], however typical empirical data does not closely realize those, and in addition the multi-scale nature of the *MSCCE* makes this derivation difficult. Another approach for normalization would be based on using a formal relationship between the *MCI* and the Spearman coefficient for Gaussian distributed vectors, for example, in analogy to the way the mutual information is rescaled when defining the *IC* [18,19]; however, this is a difficult calculation because of the complexity on estimating the entropy of the symmetrized cumulative copulas at multiple scales even in the simplest case of Gaussian distributed vectors. The exact scaling model will not have a simple closed form. As an alternative we take a more heuristic approach and use an exponential power polynomial fitted to empirical data, to numerically make the *MSCCE* approximate the Spearman correlation and in this way converge onto the interval [-1, 1] (more details on this in the next section),

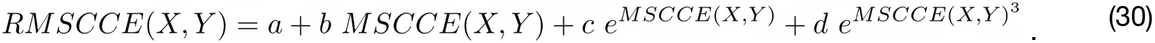

This is the final form of the *Rescaled Multi-scale Symmetrized Cumulative Copula Entropy* (*RMSCCE*) which we will refer to as the *Multi-scale Copula Index* (*MCI*). In the next section we will describe in more detail how we compute the empirical *MCI* for input data, we will also introduce a visualization paradigm for the *MCI*, analyze its computational complexity, and describe a number of benchmarks and application examples.

### Implementation of the empirical *MCI*

In order to estimate the *MCI* based on input data, we will compute empirical cumulative copulas not assuming any specific functional form. In this way the *MCI* can measure data-driven associations in a very general way without relying on specific simplifying assumptions. This is especially important in Cancer Genomics where the variables often display a complex non-linear correlation structure and display a wide range of distributional behaviors. We implemented the empirical *MCI* using a collection of finite computational lattices to provide smoothing and reduce the amount of computational work, and detect association and multiple scales. In spite of the number of operations, shown in Fig. 5, required to estimate the *MCI*, each step can be executed very efficiently for fast execution times.

**Figure 5.**
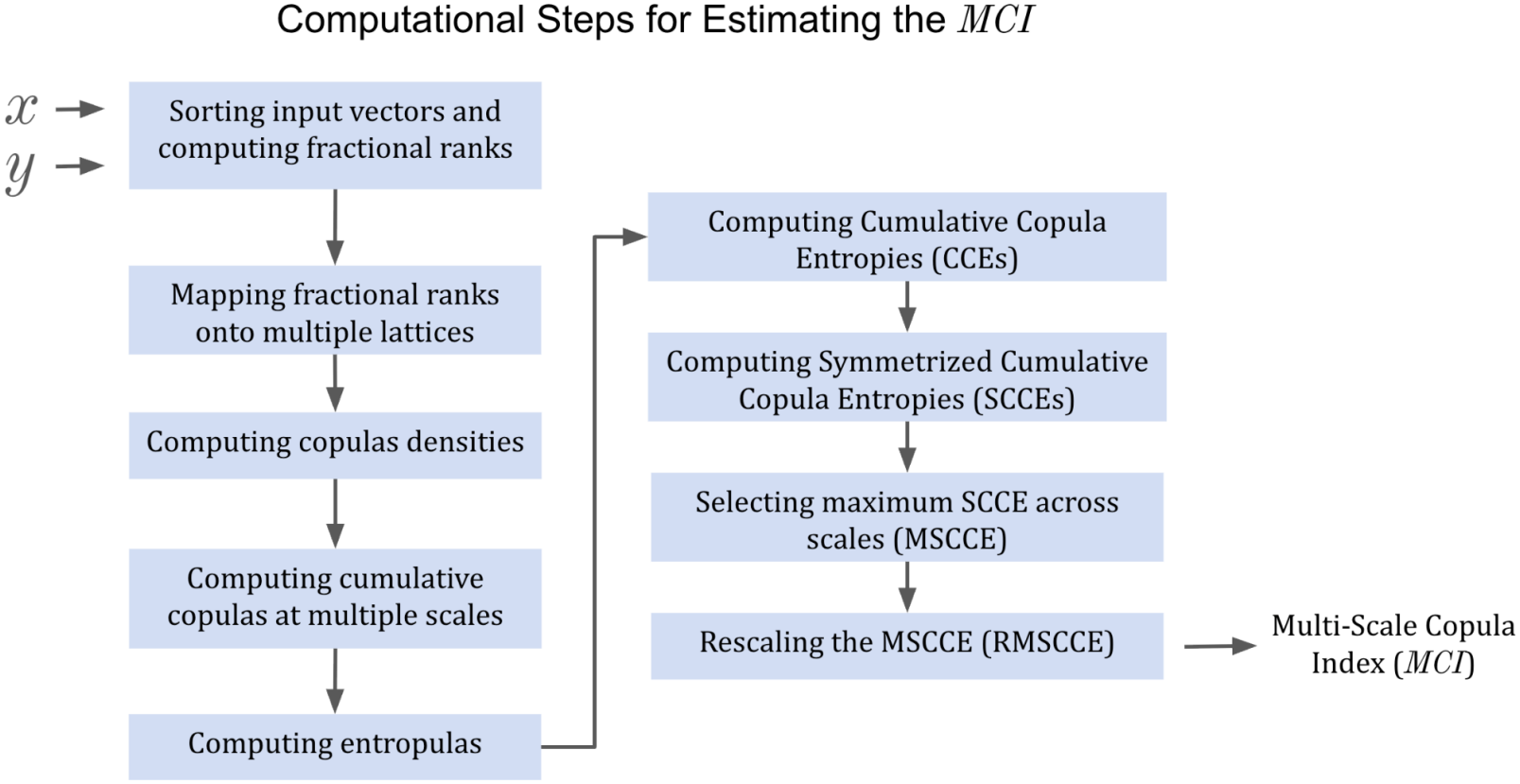
The different computational tasks involved in estimating the *MCI*.

Let’s assume we have two input vectors *X=x*_*0*_, *x*_*1*_,*…x*_*i*_,…*x*_*n-1*_, and *Y = y*_*0*_, *y*_*1*_, *…y*_*i*_,…*y*_*n-1*_, of size *n* and we are interested in computing their degree of association (see Fig. 6A). The first step in the computation is to compute their fractional ranks, starting with the ranks of *X, R*^*x*^_*i*_, and *Y, R*^*y*^_*i*_ provided by a sorting algorithm,

**Figure 6.**
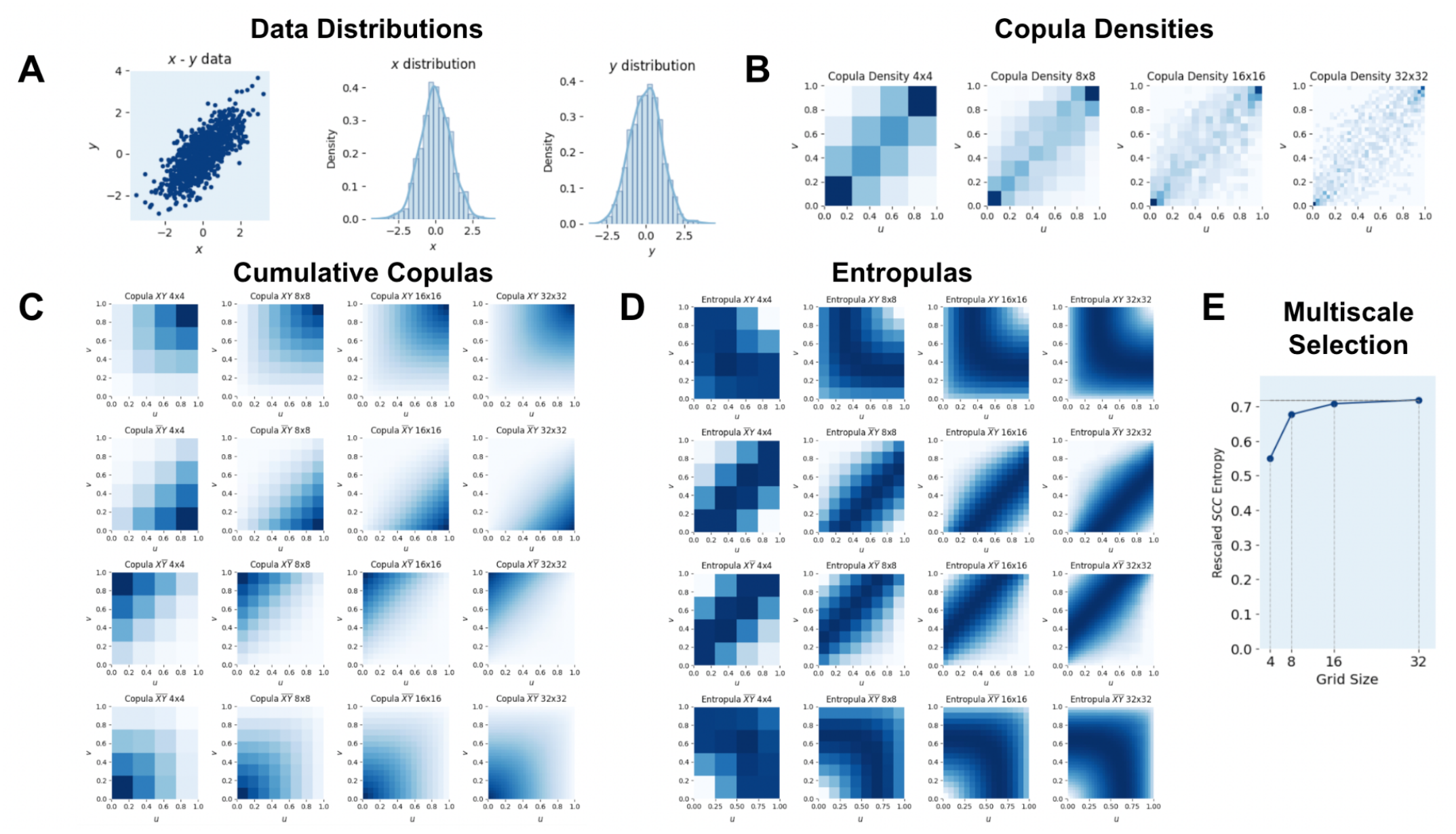
Gaussian vectors example: A) original *x* and *y* input data (1,000 points, *ϱ* = 0.85), B) copula densities, C) cumulative copulas, D) entropulas, and final selection of the maximal RSCC entropies across multiple grid sizes.

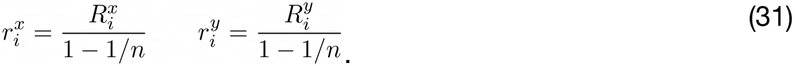

The minimum rank is zero and the maximum rank is *n-1*, and therefore the minimum fractional rank is *0* and the maximum fractional rank is 1. We can map the data points onto lattice coordinates for a group of square lattices of dimensions *g*_*min*_,…,*g*_*k*_,…,*g*_*max*_, by multiplying the fractional rank of each data point (*r*^*x*^ _*i*_, *r*^*y*^_*i*_) times the size of the lattice and flooring the results to the nearest integer,

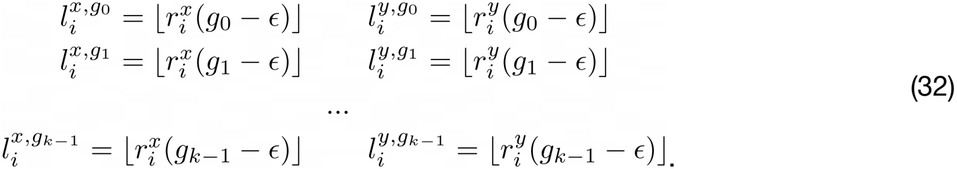

In these equations *є* is a small number (0.000001) subtracted from the lattice to keep the lattice coordinates in the range [*0, g*_*k*_ *-1*]. In this way each lattice will contain *g*_*k*_ x *g*_*k*_ points each containing the normalized count (density) of how many data points have a fractional rank corresponding to that location at the given scale of the lattice. Using these lattice coordinates for each data point (*l*^*x*^ _*r*_, *l*^*y*^_*r*_) we can compute the copula densities on the lattices at each scale,^5^ (Fig. 6B),

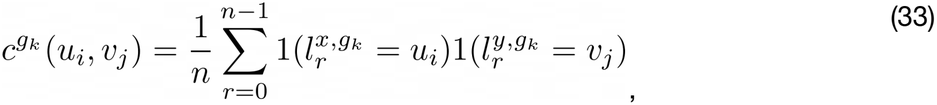

where the sample index *r* runs from *0* to *n-1*, and the lattice indices *u*_*i*_ and *v*_*j*_ run from 0 to *g*_*k*_ - *1*.

The cumulative copulas (eq. 20, Fig. 6C) can be computed by cumulative sums over the copula density at every scale,

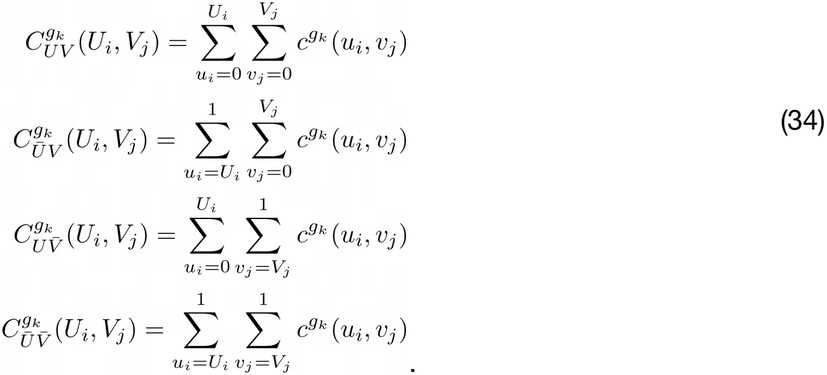

All cumulative copulas can be estimated at the same time by reindexing the lattice grids based on the desired directionality. The corresponding *entropulas* (copula entropies) for each scale *k* are generated from the cumulative copulas (Fig. 6D),

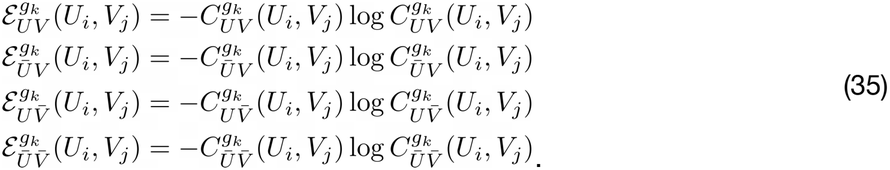

The cumulative copula entropies (*CCE*) are then,

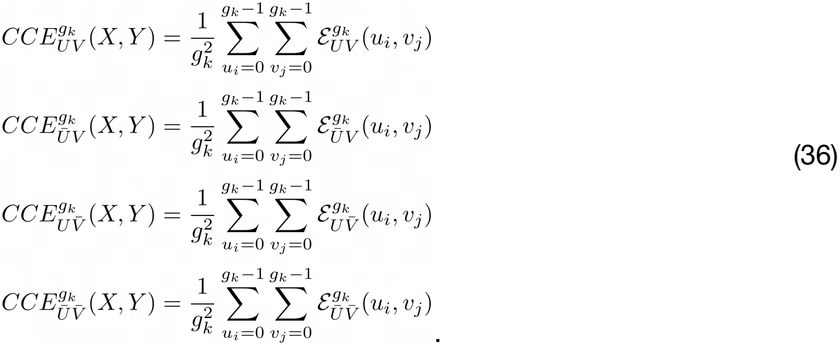

The symmetrized version of the cumulative copula entropies (*SCCE*) for each scale are obtained by an additive combination of the direct and survival copulas then subtracting both hybrid ones,

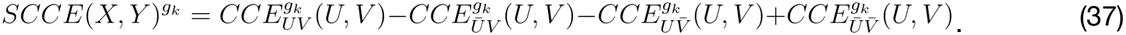

The multi-scale, symmetrized, cumulative copula entropy (*MSCCE*) is the maximum across the different scales (eq. 29, Fig. 6E),

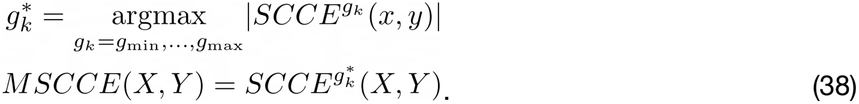

Finally, the rescaled *MSCCE* (*RMSCCE*) is obtained by rescaling the *MSCCE* using a heuristic^6^ 4-parameter exponential power polynomial in order to make it approximate the Spearman coefficient for Gaussian-distributed vectors,

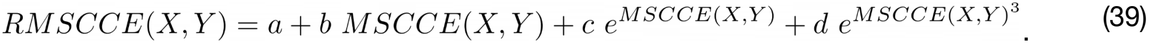

This rescaling makes the resulting *RMSCCE* behave like the Spearman coefficient when the variables have a linear relationship. In Fig. 7 we show the fitted 4-parameter model using Gaussian distributed vectors of size 1,000 and variable correlation from -1 to 1 (the figure shows only the positive part). As can be seen in the figure, this scaling model makes the *MCI* approximate the Spearman coefficient for Gaussian distributed vectors extremely well. Because the vector size dependence of the MCI is very small (Fig. 9) this normalization is used generally regardless of the size of the input vectors. To avoid the long acronym we rename the *RMSCCE* as the *Multi-scale Copula Index (MCI)* for short.

**Figure 7.**
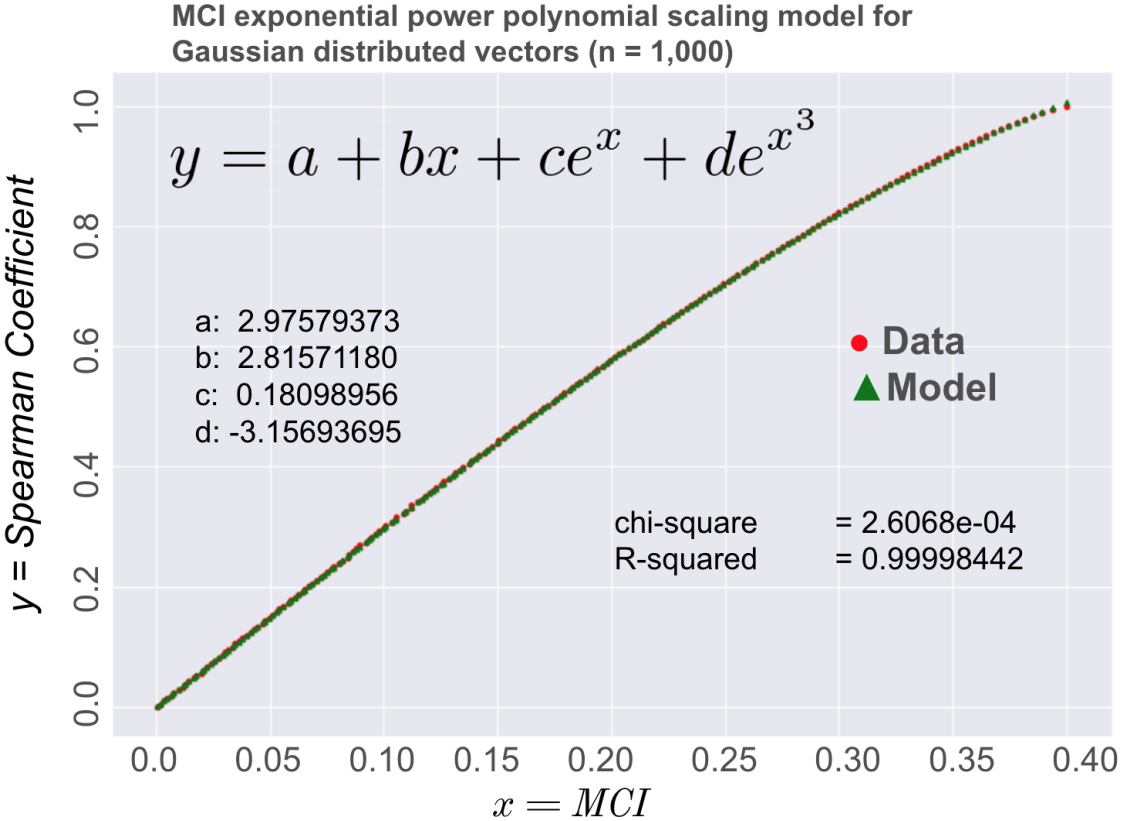
Empirical scaling model for the *MCI:* the 4-parameter exponential power polynomial model is fitted to make the *MCI* approximate the Spearman correlation for Gaussian-distributed vectors (n = 1,000).

**Figure 8.**
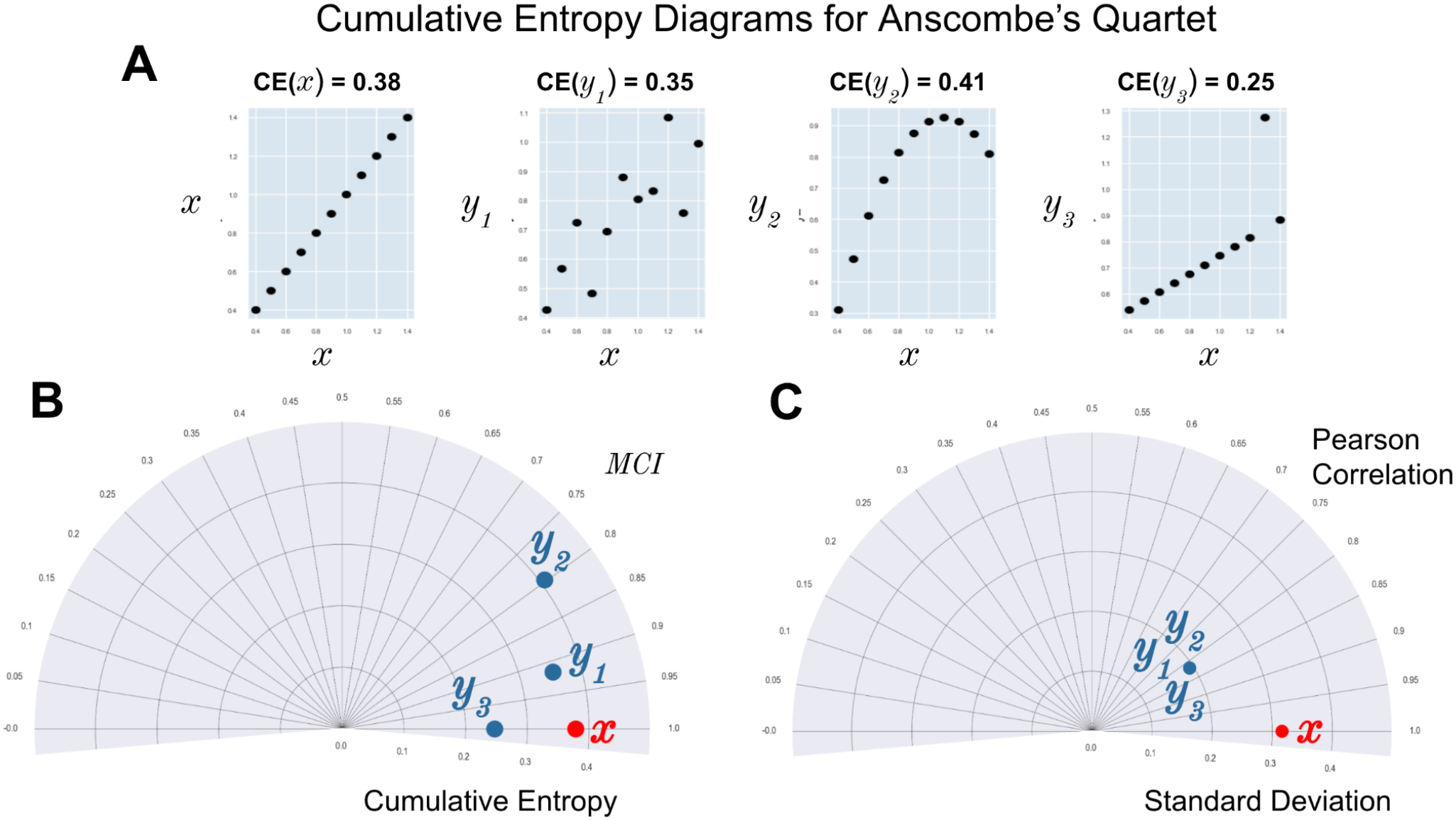
Data distributions for Anscombe’s quartet (A), cumulative entropy diagram showing the *MCI* and Cumulative Entropy (B), same diagram but using the Pearson correlation and standard deviation.

**Figure 9.**
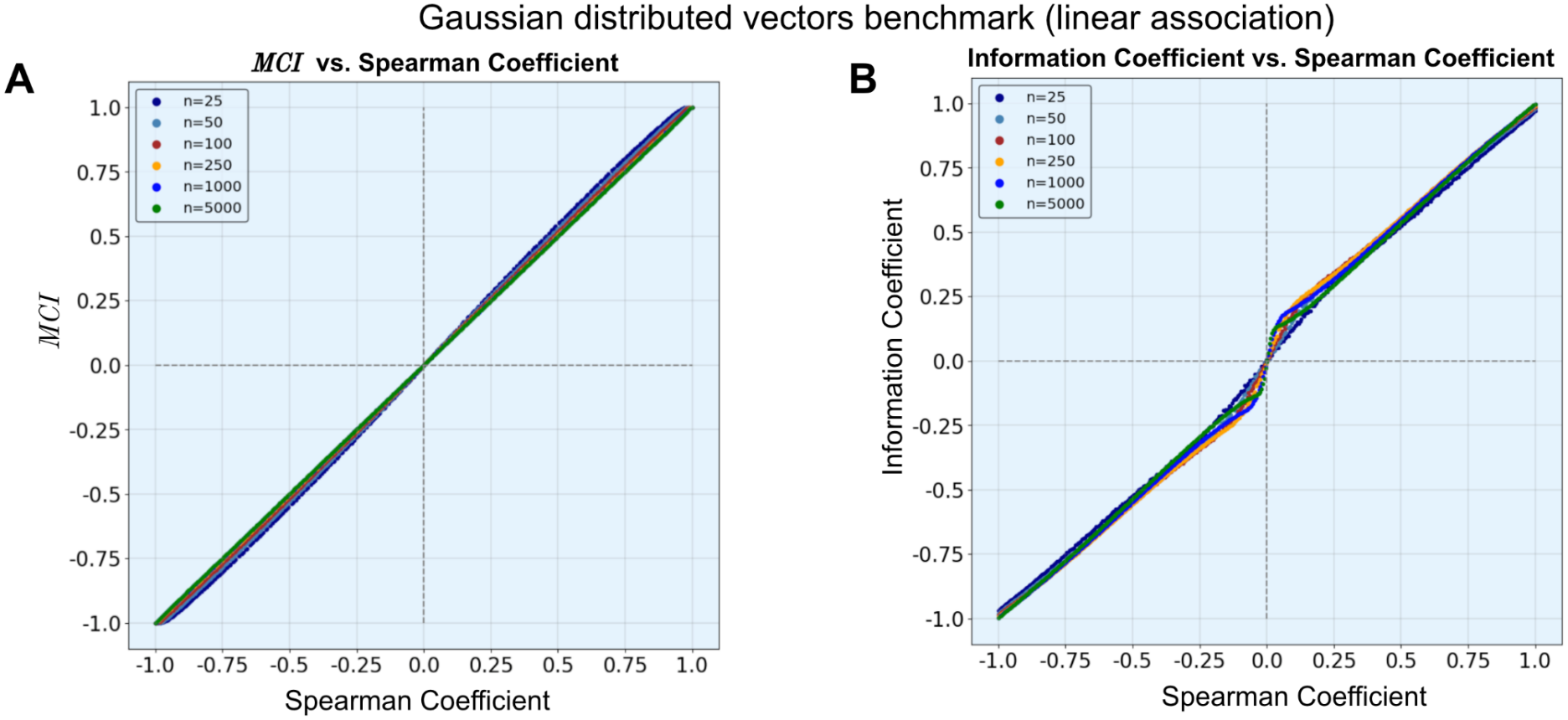
Gaussian-distributed data benchmarks for A) the *MCI* and B) the Information Coefficient (*IC*).

At the end we include a link to the GitHub repository that hosts a copy of the computer code that implements the *MCI* as described above. This version has been customized to serve as a measure of association in our Cancer Genomics applications described below. The implementation requires us to make choices about the number of scales which we choose to be power-of-two lattices from 4 to 32: *g*_*4×4*_, *g*_*8×8*_, *g*_*16×16*_, *and g*_*32×32*_. The use of compiled Cython code, to implement the computation of the copula density, the cumulative copulas and the entropies, makes the implementation of the *MCI* over 30 times faster than our kernel-based implementation of the mutual information/Information Coefficient (IC).^7^

### Computational Complexity of the *MCI*

Assuming *n* is the size of the input vectors (*x, y*) and *k* is the size of the largest grid. The main computations of the *MCI* computation, and their corresponding approximated computational complexities, are: i) an argument sort (*argsort*) for each *x* and *y* variable: *O*(*n log n*), ii) the computation of the fractional ranks for each variable: *O*(*n*), iii) the frequency computation to compute the copula density: *O*(*n*), iv) the computation of the cumulative copulas: *O*(*k*^*2*^), and, v) the computation of the entropies: *O*(*k*^*2*^). As the grid size (*k*) is typically much less than the input vector length (*n*), then the computation of the *MCI* would be dominated by the *O*(*n log n*) *argsort* scaling.

### The case of binary vs. continuous inputs to the *MCI*

When one of the input variables is binary rather than continuous, the standard probability integral transform does not result in a continuous Uniform (0, 1) distribution. Because a discrete random variable only takes on specific, separated values, its cumulative distribution function (CDF), *F*(*X*) is a step function and consequently, applying the CDF to the random variable itself results in a new variable that is also discrete but fails to be uniform. In this case, to recover a continuous Uniform(0, 1) distribution from a discrete random variable, we use a technique called the randomized probability integral transform, sometimes also called continuous extension or jittering. The idea behind this approach is to “fill in the gaps” of the step function by adding a continuous, uniform random noise between the steps. The implementation of the MCI, when one of the input variables is binary, makes use of a Randomized Probability Integral Transform [52] where random Gaussian distributed noise is added to the binary variable in order to break the ties and allow the resulting variable be able to be mapped in a continuous Uniform (0, 1) distribution. In example 5 we show two examples of matching genomic data that use the *MCI* in the case where one of the variables is binary.

### Cumulative Entropy Diagram (CED)

In order to provide a visual paradigm that allows the visualization of the *MCI* between a reference vector and multiple feature vectors we introduce here the *Cumulative Entropy Diagram* (*CED*). The *CED* was inspired by the *Mutual Information Diagram* (*MID*), introduced by Correa and Lindstrom [53], which in turn was motivated by the classic Taylor Diagram [54] of common use in climate science, meteorology, and environmental modeling. The Mutual Information Diagram is used to quantify and visualize the association and statistical dependence of two or more variables. It measures how much information one variable provides about another, based on their mutual information and it is useful to visualize both linear and non-linear relationships. The Taylor Diagram provides a graphical summary of how well a model or set of predictions matches observations. It simultaneously represents the correlation, standard deviation, and root mean square error (RMSE) of a dataset. The most straightforward approach to generate an *informational diagram* for the *MCI*, would be to exploit the relationship between the marginal and joint entropies and generate a geometric relationship between these using the law of cosines [53]. However, because the *MCI* has no simple relationship with the marginals’ cumulative entropies, we do not rely on the law of cosines and just plot the angular distance proportional to the *MCI* and the radial distance proportional to the cumulative entropy of the marginal variables as shown in Fig. 8. In this way the *Cumulative Entropy Diagram* (CED) helps to visualize the relationships between a reference variable and a group of additional variables, and their intrinsic cumulative entropies. This is illustrated for the Anscombe’s quartet^8^ dataset in Fig. 8A where one can notice how the variables *y*_*1*_, *y*_*2*_ and *y*_*3*_ are clearly distinguished using the *MCI* and Cumulative Entropies *CED* (Fig. 8B), but not when using the Pearson correlation and the standard deviation where all 3 points are superimposed on each other in the diagram (Fig. 8C). Other examples of *CED* diagrams for real genomics data are shown in the applications section.

### Generalizing the MCI to higher dimensions

Extending the *MCI* to higher dimensions is conceptually straightforward because Sklar’s theorem and the copula entropy relationship, already generalize to *n-*dimensional multivariate distributions, as shown in eq. 9, which defines the copula for *n* variables *X*_*1*_,…,*X1*. The empirical implementation would require extending the computational lattices from 2D grids (*g*×*g*) to *n-*dimensional hypercubes (*g*×*g*×…×*g*), computing *n-*dimensional cumulative copulas via multiway cumulative sums, and generalizing the symmetrization scheme (eq. 24) to account for all 2^n^ combinations of forward/backward integration directions across each variable’s axis, analogous to how the bivariate case used 4 combinations: direct, survival, and two hybrid copulas. This would allow one to define a Multi-scale Copula Interaction Information (MCII), in a similar way how classical interaction information for three or more variables is built from alternating sums of joint and marginal mutual informations/entropies [55,56], in this case instead built from alternating sums of the multi-scale symmetrized cumulative copula entropies for all subsets of the variables. Such a measure would quantify higher-order, non-additive dependencies among three or more genomic variables, e.g., synergistic or redundant relationships between a gene’s mRNA, CRISPR sensitivity, and a drug response phenotype, that cannot be captured by pairwise *MCI* alone, while retaining the measure’s key practical advantages: no density estimation, directionality, robustness, and computational tractability relative to kernel-based multivariate information measures. The main challenges estimating this measure are related with the combinatorial growth in the number of hybrid copulas and lattice sites with increasing dimension *n*, requiring careful attention to computational complexity and sample size requirements in higher-order lattices.

### *MCI* benchmarks

Here we study the performance of the *MCI* in terms of the quality of the association as well as its execution times. These provide evidence of the power of the *MCI* measure to detect statistical dependency and overall associations and its practical utility and computational properties.

#### 1. Gaussian-distributed synthetic data

To study the behavior of the *MCI* across a wide range of association strengths we generated gaussian-distributed vectors (*x, y*) of length *n* = 10, 25, 50, 100, 250, 1,000, and 5,000 with predefined standard correlation *ϱ*(*x, y*), marginals means equal to 0, 1 and marginal standard deviations equal to ½ and ½ ^9^. We generated synthetic data instances starting at *ϱ*(*x, y*) = -1 (complete anti-correlation), and up to *ϱ*(*x, y*) = 1 (perfectly correlated) and all intermediate correlations at intervals of 0.01. The vectors were truncated above and below 3 standard deviations to avoid large value outliers, and the benchmark was iterated, by generating different randomly-generated instances with the same correlation, and averaged over 2,500 times for each vector size (*n*).

Fig. 9 shows the results using the *MCI* (eq. 35) and the *IC* (eq. 6) measures to assess the degree of association of Gaussian-distributed vectors. The graphs show the expected behavior, i.e., the proportionality between the *MCI*(*x, y*) and the *IC*(*x, y*) versus the correlation *ϱ*(*x, y*) between pairs of vectors. Compared with the *IC*, which is a rescaled mutual information, the *MCI* shows two remarkable characteristics: it shows practically no finite-size effects, as it is essentially the same between two vectors having the same correlation regardless of the vector size (Fig. 9A). In contrast, the *IC* shows some degree of finite-size variation as a function of vector size, particularly at low values of *n* (Fig. 9B). Another interesting characteristic is the smoothness of the *MCI* transition around the zero correlation point compared with the more sudden and steeper change in the *IC*. These two behaviors for the *IC* measure, i.e., higher finite-size variation and discontinuous zero-correlation crossing, are both produced by limitations in the estimation of the kernel density bandwidth necessary to compute the *IC*. Even though the *x* and *y* bandwidths have been adjusted to take into account the correlation in our implementation of the *IC*, this correction is not enough to eliminate those effects. This was one of the main motivations for developing the *MCI* as it requires no estimation of kernel densities and their associated bandwidths.

#### 2. Synthetic data: simple and complex dependencies benchmarks

In these benchmarks we generated synthetic data to represent different types of pre-defined associations between two random variables and used the *MCI* to quantify the degree of this association. The associations include, for example, *linear, polynomial* and *hyperbolic tangent* functions as shown in Fig. 10 below. For each type of association we generate several versions of the *x* and *y* random variables by changing one parameter (*a*) of the functional form and in this way modulate the amount of correlation between the variables. We also add some amount of uniform random noise (*b*) to each instance of the benchmark in order to study the robustness of the association measure. For every variation of the benchmark we generate 1,000 data points and measure the degree of association using the *MCI* and the Spearman coefficient and average over 500 instances using different random numbers.

**Figure 10.**
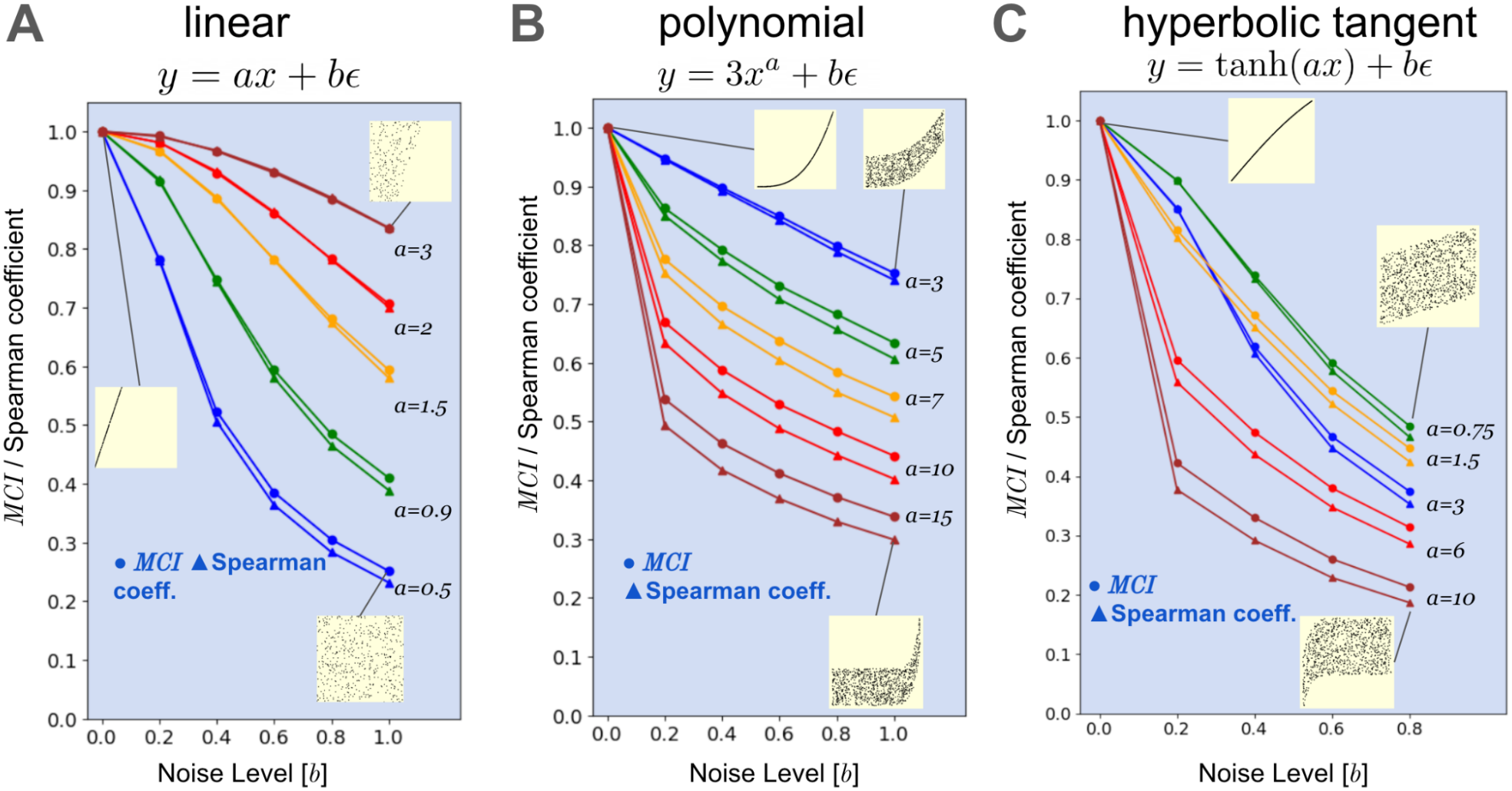
Benchmarks results: simple functional associations: A) linear, B) polynomial and C) hyperbolic tangent.

The benchmark results are shown in Fig. 10 using circles for the *MCI* and triangles for the Spearman coefficient. In the linear benchmark (Fig. 10A), the differences between the *MCI* and the Spearman coefficient, are small when the noise is small but become more visible for larger amounts of noise. This shows that there is an advantage of using the *MCI* even for variables that show a linear relationship but may have some amount of noise (Fig. 10A). The differences are more noticeable for the polynomial and hyperbolic tangent benchmarks (Fig. 10B-C).

Fig. 11 shows a set of 3 more complex functional dependencies, including *sine power, scaled logistic* and *lizard-shaped*. In all these cases the benchmark results show the advantages of the *MCI* in terms of i) higher sensitivity to the association signal, and ii) increased robustness to noise. These two properties make the *MCI* specially valuable to be used in functional cancer genomics when comparing variables with different types of statistical association as we will show in the examples and applications section.

**Figure 11.**
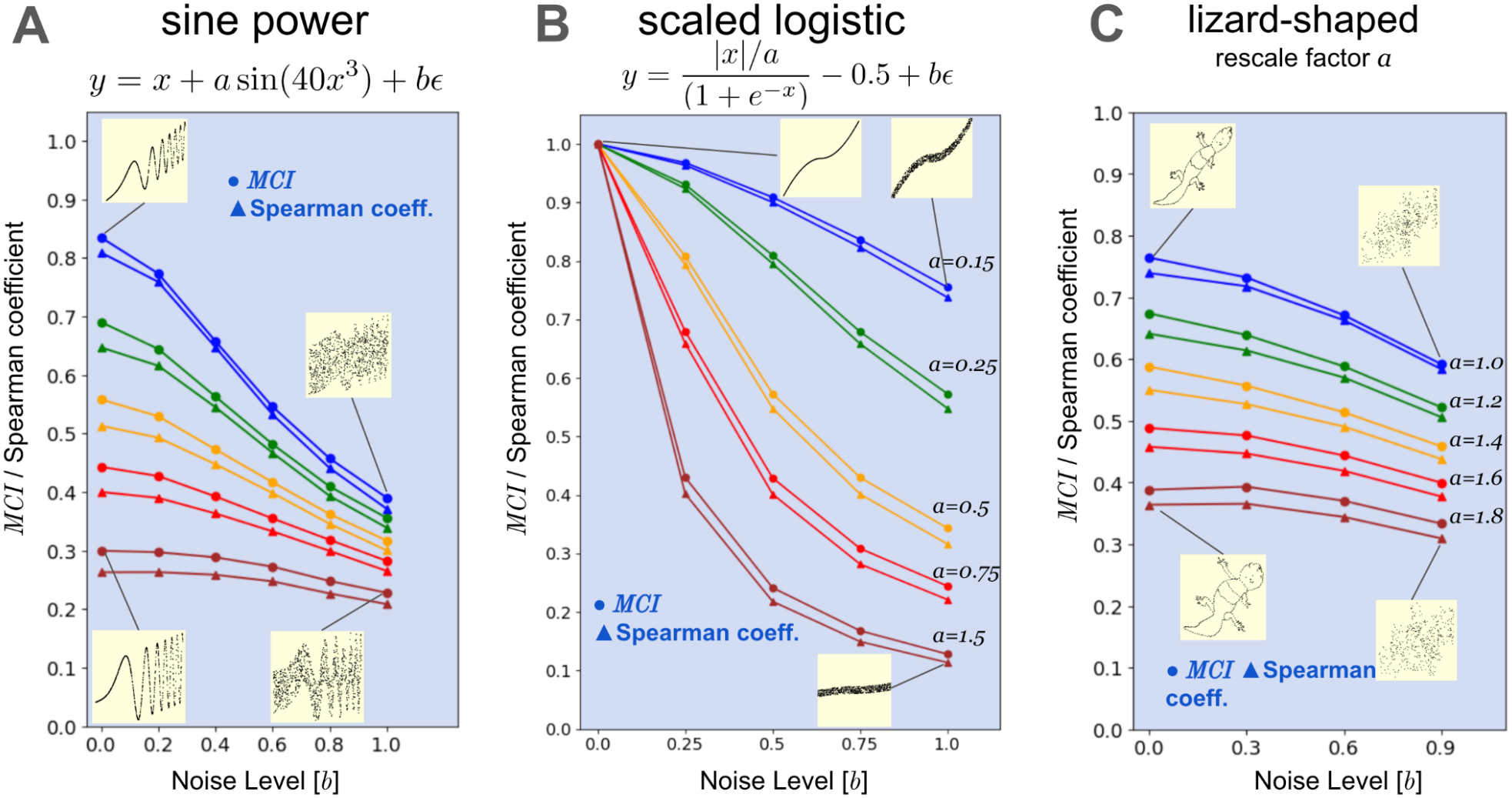
Benchmarks results: more complex functional associations: A) sine power, B) scaled logistic and C) lizard shaped.

#### 3. Timing benchmarks: *MCI* vs. Spearman correlation and the *IC*

We performed execution time benchmarks to analyze the running time performance of the *MCI* and compared against the Spearman correlation and the *IC* for the same Gaussian input vectors varying in size from 200 to 2,000 data points and averaging the execution times over 1,000 instances. The timing results^10^ are shown in Fig. 12, where we can see that the *MCI* is about 35 times faster than the *IC*, and comparable to the Pearson coefficient computation for vectors of less than 1,000 data points. This a significant increase in performance over the kernel density-based *IC* and opens the way to use the *MCI* to compare genome-wide molecular variables with themselves or against other datasets and systematically associate each of them with all the others. Its high performance makes the *MCI* a very useful and practical measure to generate large association matrices that in turn can be used to build molecular networks [57,58], cluster samples etc. We show an example of this type in the examples and application section below.

**Figure 12.**
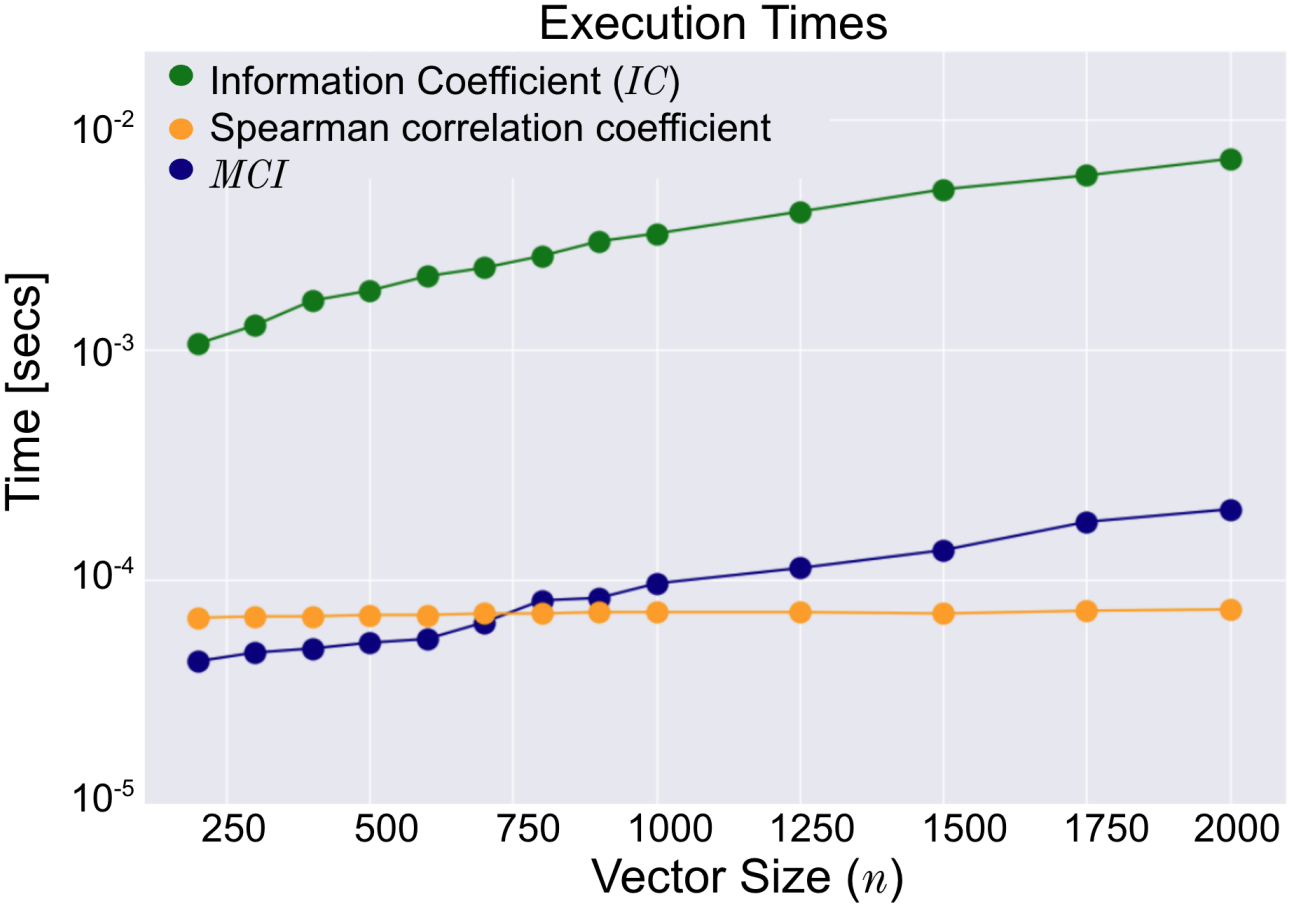
Timings for the computation of the *MCI*, Information Coefficient (*IC*) and the Spearman coefficient.

### Functional Cancer Genomics Examples and Applications

As we mentioned in the introduction, integrative cancer genomic analysis works with large and complex datasets that integrate various ‘omics data types [1–5]. In this context, it is important to be able to compare diverse genomic variables to establish associations and select candidates for further study or validation. However, these comparisons require caution due to the complex correlative structure of genomic data. Despite their importance, current feature selection and association methods often use basic measures and algorithms that make overly simplistic assumptions about the data’s statistical properties. This is particularly relevant when using perturbational, drug sensitivity or functional genomic data such as those generated over the last few years to measure the effects of a large number of genetic or chemical perturbations on transcription or cell viability. These include e.g. the Cancer Cell Line Encyclopedia (CCLE) [59], The Cancer Dependencies Map [23,60], the Cancer Therapeutics Response Portal [61–63], the Genomics of Drug Sensitivity in Cancer [64–66], the Library of Integrated Network-based Cellular Signatures (LINCS) [67], and the Cancer Cell Line Factory project [68,69]. Many of these datasets are now consolidated into the Dependencies Map website (depmap.org) at the Broad Institute, or are part of the *Cancer Target Discovery and Development Network* (CTD^2^) [70], a functional genomics initiative supported by the National Cancer Institute. Some of those efforts have focused on systematically identifying genome-wide profiles of gene essentiality across large collections of genomically characterized cancer cell lines using genetic perturbations such as RNAi and CRISPR-Cas9 that silence or knockout individual genes. These datasets have become an invaluable resource, for example, to identify synthetic lethalities [71,72], essential genes downstream of oncogenic pathways [73], novel vulnerabilities associated with copy number alteration [74–76], tissue or subtype vulnerabilities [60,77], and the functional status of large groups of oncogenic alleles and genetic predictors of gene dependency [78]. These resources have allowed expansion of the diversity of therapeutic targets engendered by non-oncogene dependencies to a much larger universe than the one defined by recurrently mutated genes [6]. In the next few sections we will show the advantages of more sophisticated information-based measures of association, such as the *MCI*, that are able to measure associations between genomic variables with non-linear relationships and that do not make overly simplistic assumptions about the data distributions.

#### Example 1. Estimating the degree of association of genes (mRNA) with EGFR’s protein expression

In this example we use the protein expression profile of EGFR (RPPA^11^) as a reference phenotype and we match to it the gene expression (RNA-Seq) profiles of 16,184 genes in 893 cancer cell lines using the *MCI*. The relevant datasets were downloaded from the publicly available Broad Institute’s DepMap (depmap.org). The computation of the *MCI* takes only about 2.8 seconds. We also randomly permuted the EGFR RPPA profile 1,000 times and matched all the genes profiles against these randomized profiles as part of a permutation test [79] to estimate the statistical significance of the scores, namely, the p-values and False Discovery Rates (FDRs). The entire process involves the evaluation of 16,184,000 *MCI* associations between 893-long input vectors (RPPA and mRNA) and takes about 40 minutes in a laptop. Fig. 13 shows a heatmap of the top/bottom 25 genes most positively/negatively associated with the EGFR protein profile where the reference and genes profiles have been standardized and clipped at 2.25 standard deviations. The color mapping shows red for feature values corresponding to values above the mean, and blue for values below the mean, while white represents the mean. The top hit, namely the most associated gene (mRNA) with the EGFR protein profile is the EGFR (mRNA) itself (*MCI* = 0.90 ± 0.015). The entire distribution of *MCI* values for all genes

**Figure 13.**
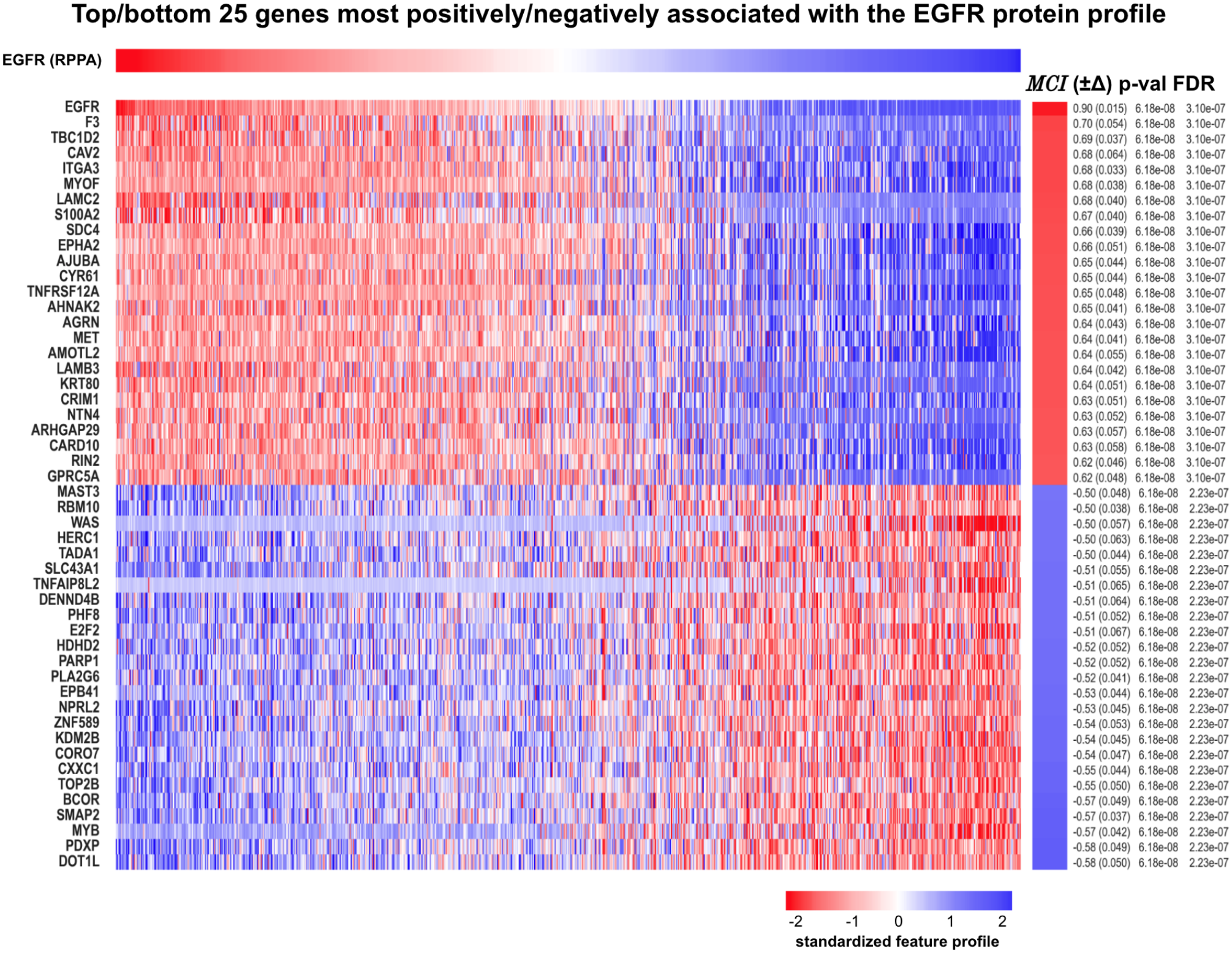
Heatmap showing the top (bottom) 25 genes most positively (negatively) associated with the reference EGFR protein profile according to the *MCI*.

**Figure 14.**
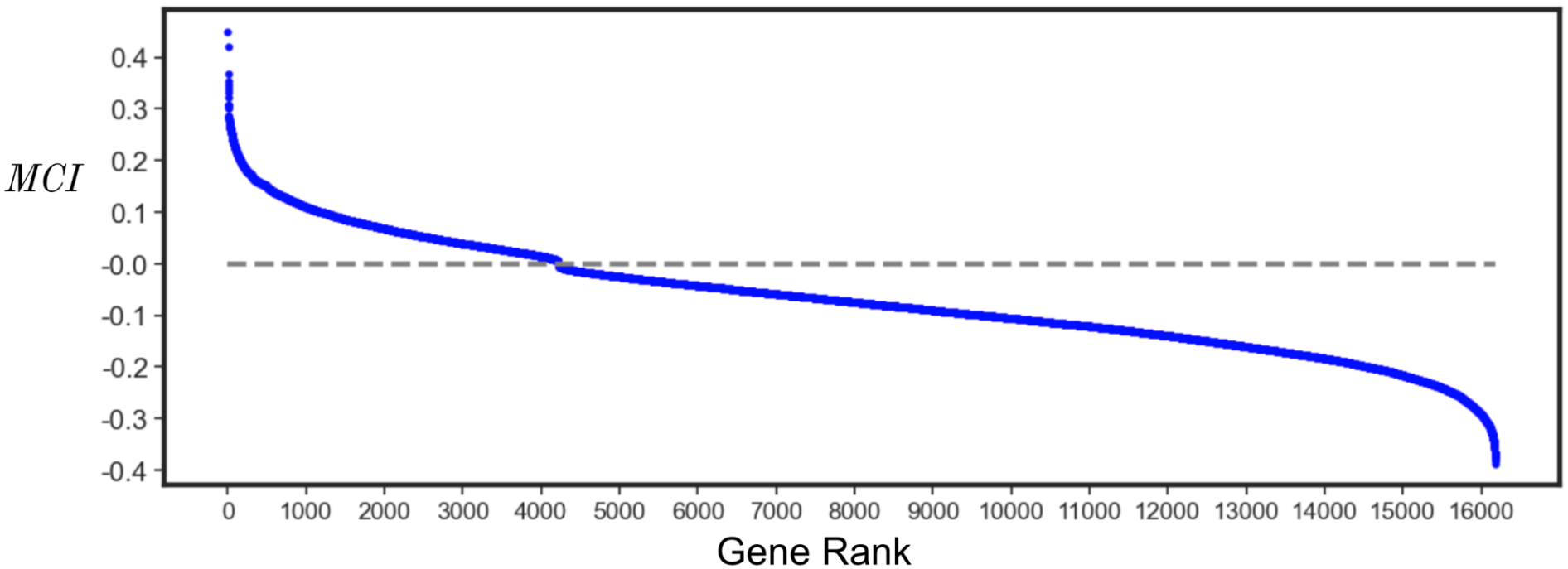
*MCI* values for all genes (16,184) sorted from maximum to minimum.

If we look more carefully at the top 50 hits we can easily identify 17 genes that have been implicated in the EGFR pathway. These genes are shown below in Fig. 15 using the *Cumulative Entropy Diagram* (*CED*). In this diagram, the blue circles correspond to those 17 genes, and the red circle shows the EGFR (protein) reference profile. The orange circles represent the other 16,167 genes in the dataset where the size of the circle is proportional to the *MCI*. The radial direction in the diagram corresponds to the cumulative entropy of each gene, with higher radii representing features with higher cumulative entropy. The angular direction corresponds to the *MCI* between each gene and the reference with higher values to the right side of the diagram. In this diagram one can clearly see that, as we noticed before in the heatmap, that the EGFR gene (mRNA), has a much higher association against the reference than the other genes, and attains an *MCI* of 0.90, significantly higher than the other genes which have *MCI*s in the 0.50-0.70 range. This gap is much easier to see in the *CED* than in the heatmap. In the diagram we can also see that the cumulative entropy of genes such as ITGA2 and MET is less than all the other genes and that MYOF attains the highest cumulative entropy. We also see that the EGFR gene (mRNA) profile has somewhat higher cumulative entropy than the reference EGFR protein profile. This *CED* example shows the utility of the diagram to analyze many associations against a reference profile as it is often the case in functional genomics analysis. The *CED* can also be parametrized to show the entire 360 degrees layout and display negative associations in the lower part (not shown).

**Figure 15.**
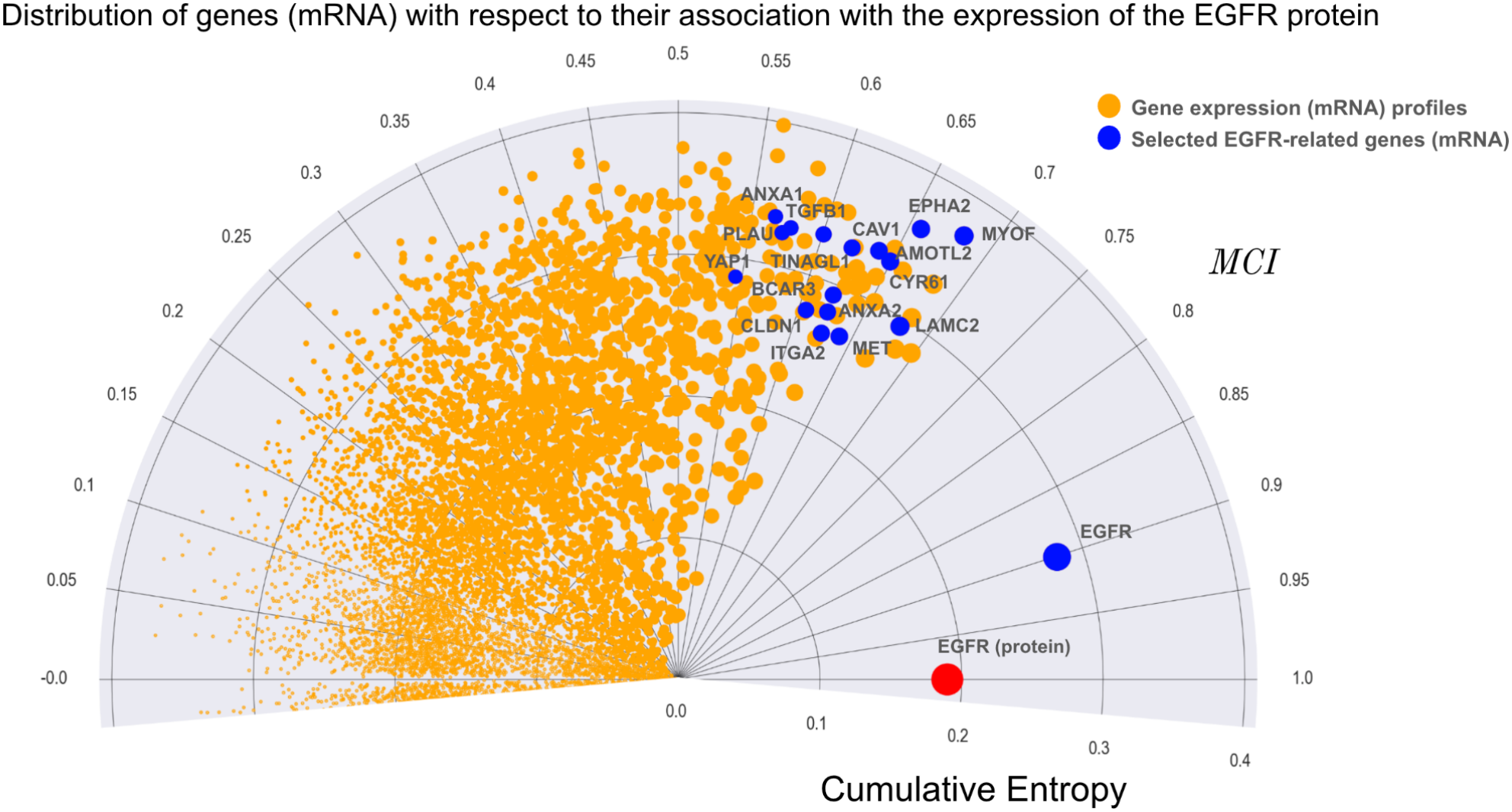
Cumulative entropy diagram showing all the mRNA profiles (orange) with respect to the reference EGFR protein profile (red). A subset of 17 EGFR-related genes are highlighted in blue.

#### Example 2. Associating transcriptional and perturbational profiles of oncogenes

Here we analyze in more detail the example used in the introduction on the statistical dependency between mRNA and CRISPR/Cas9 profiles for the same oncogene in large numbers of cancer cell lines. As we described in the introduction the *MCI* measure was able to better detect the association of PAX8 CRISPR and mRNA profiles. This behavior is also seen in the other examples as shown in Fig. 16 where we compared the association values according to the *MCI* and the Spearman coefficient for the 12 genes. The *MCI* measures (blue bars) are all statistically significant with p-values below 10^6^ for those genes.^12^ We can see in the figure that all the *MCI* values (blue bars) are higher than Spearman’s (orange bars) suggesting that the mRNA-CRISPR relationships for all those oncogenes are at least partially non-linear. This type of contextual association, when two genomic variables are uncorrelated for most of the data points except for a subset representing the “active” biological context where there is a strong association, is common in functional genomics readouts and therefore an opportunity to use the *MCI* to quantify those associations.

**Figure 16.**
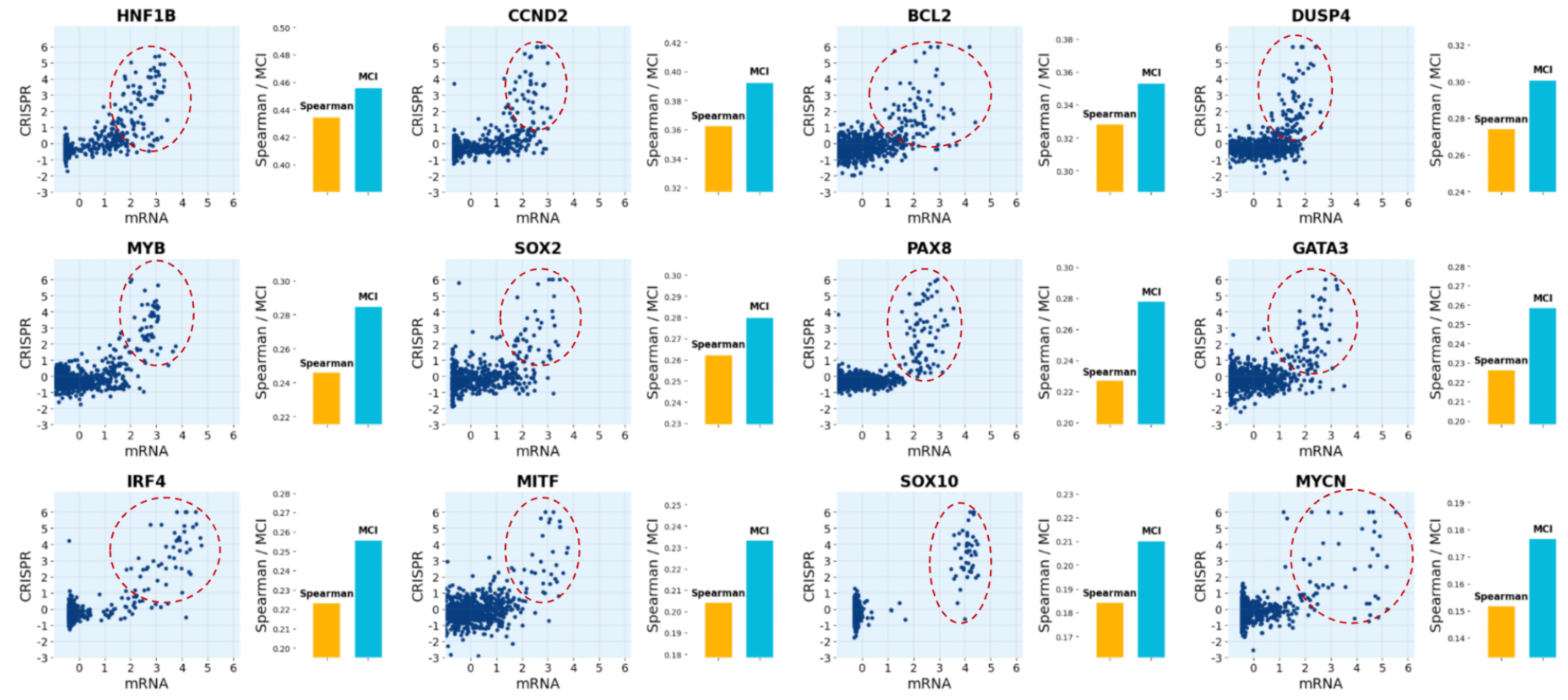
Examples of relationships between CRISPR and mRNA profiles for 10 selected oncogenes and the corresponding Spearman correlation and *MCI* quantitative measures of association.

A more detailed analysis of the association between the mRNA and CRISPR values for the MYB oncogene is shown in Fig. 17 where we can see the histograms and fitted 4-parameter *skew-t* distributions [80] for MYB’s individual mRNA and CRISPR distributions. The extreme values, higher expression and higher CRISPR sensitivity produce very long non-exponential tails. The association is much stronger in a specific context, presumably a cellular state or states, where the oncogene is active or significantly induced and therefore the cancer cells are more dependent on it. In other contexts, the association is much weaker and not particularly significant, e.g., in the bulk of the distributions.

**Figure 17.**
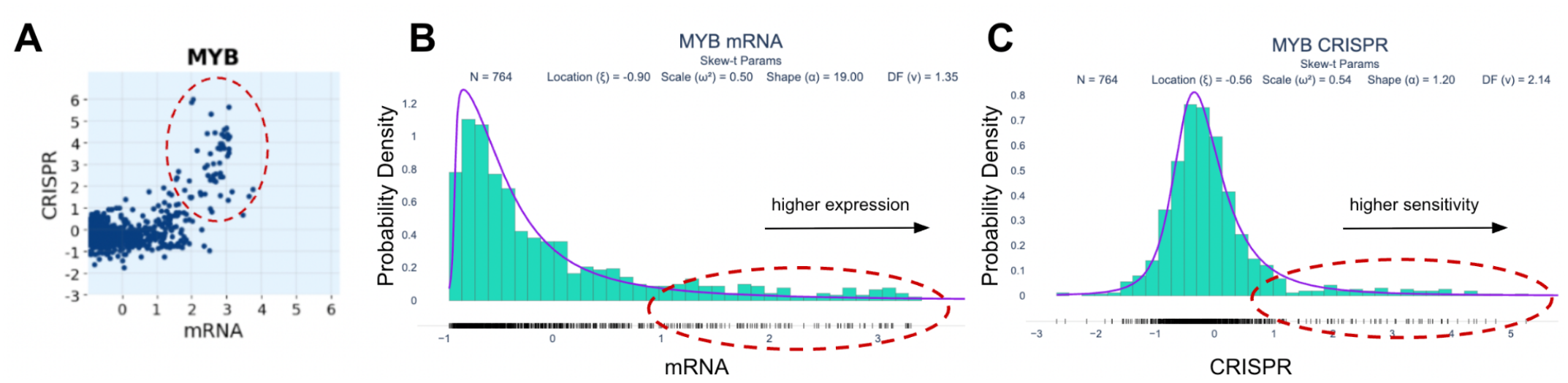
A) Relationship between MYB (oncogene) mRNA profile (x-axis) versus the negative of its CRISPR/Cas9 cell viability profile. B-C) Histograms and 4-parameter *skew-t* fits of the MYB mRNA and CRISPR distributions.

We will also use this example to illustrate the multi-scale nature of the *MCI*. In Fig. 18, we show examples of mRNA-CRISPR relationships at the level of the multiple scales’ symmetrized cumulative copula entropies (SCCE), where the maximum SCCE, used to define the final (maximum) value that defines the *MCI* takes place at one of different grid sizes such (4 × 4, 8 × 8, 16 × 16 and 32 × 32). Often the highest value takes place at the finest scale (32 × 32) but this is not always as can be seen in the figure.

**Figure 18.**
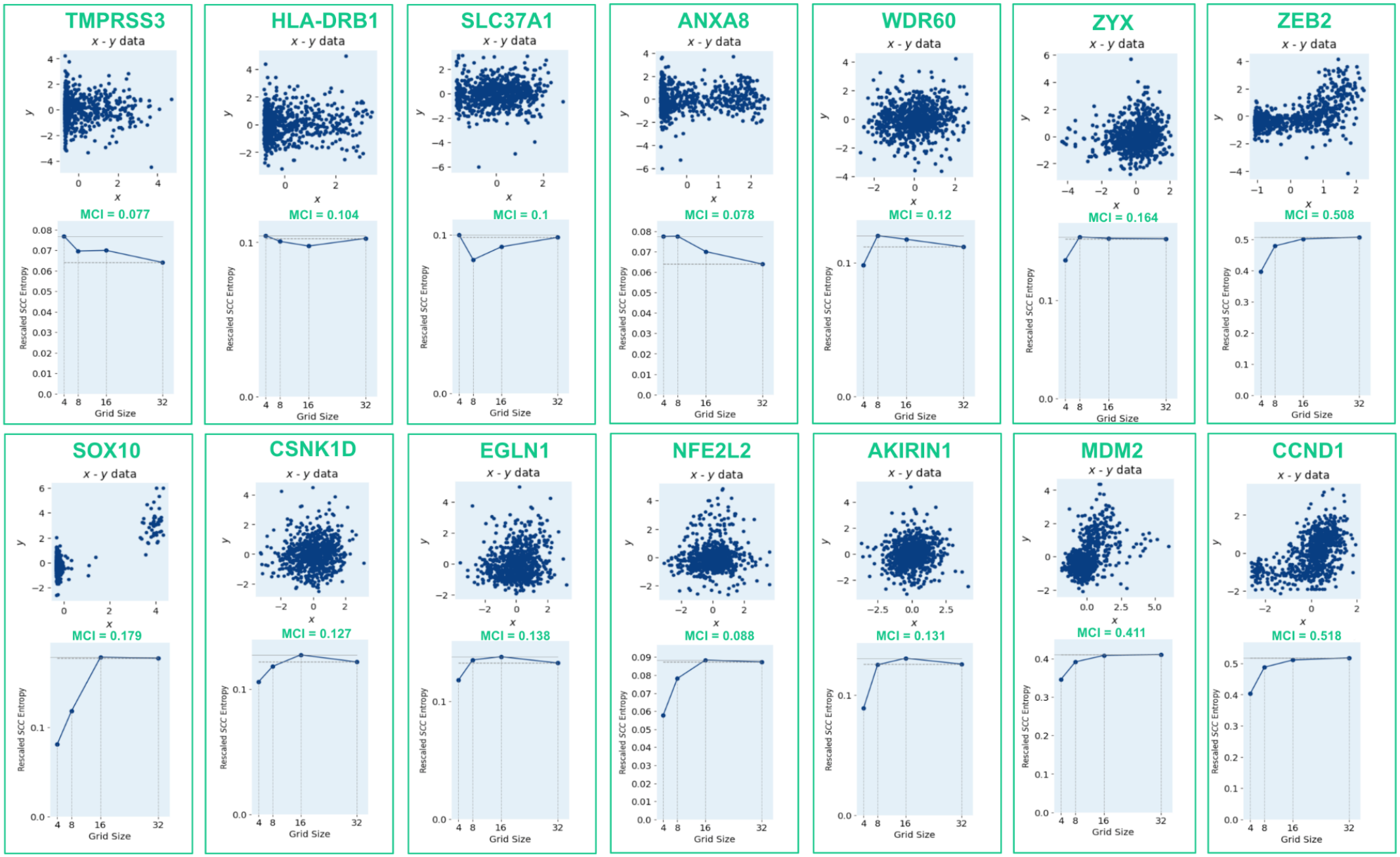
Examples of mRNA-CRISPR gene relationships where the largest *MCI* peaks at different scales, i.e., grid sizes (4×4, 8×8, 16×16 and 32×32).

#### Example 3: generating a trametinib drug response signature

In this example we used the *MCI* to find genes associated with the Broad Institute’s PRISM drug-screen (PMID: 26928769) response to Trametinib, a targeted cancer drug that blocks specific proteins (MEK1 and MEK2) involved in cell growth and division pathways that are often overactive in cancer cells. This drug is primarily used to treat advanced melanoma, particularly in patients whose tumors have BRAF gene mutations, and is frequently combined with other targeted therapies like dabrafenib for enhanced effectiveness. The drug is also approved for treating certain other cancers with specific genetic alterations, including some thyroid cancers and solid tumors with BRAF mutations. To perform this analysis we used the CCLE mRNA dataset and we chose the subset of cancer cell lines for which the response to trametinib had been measured as part of the PRISM screen. The Broad Institute’s PRISM (Profiling Relative Inhibition Simultaneously in Mixtures [81] is a high-throughput drug screening platform that tests thousands of compounds against hundreds of cancer cell lines simultaneously using DNA barcoding technology. As part of the Cancer Dependencies Map (DepMap) project, this screening effort has generated comprehensive dose-response data to identify which cancer cell lines are sensitive or resistant to specific drugs, helping researchers discover new therapeutic vulnerabilities and understand drug mechanisms of action.

To generate a drug sensitivity vector we used the negative of the PRISM relative readout of cell number for trametinib. In this way we were able to generate a vector of sensitivities and use the *MCI* to find genes that have expression profiles that match that sensitivity vector. First we split the CCLE mRNA and PRISM drug response overlapping datasets (466 samples) into train and test subsets containing 308 and 158 samples respectively. Then we used the train set to sort genes (mRNA) according to their association to the trametinib drug response vector using the *MCI* (Fig. 19). We performed an empirical permutation test and repeated the entire computation for 1,000 random permuted versions of the drug response profile and then used the original and these permutation test *MCI* readouts to compute p-values and False Discovery Rates (FDR). These measures of statistical significance are also shown on the right side of the heatmap in Fig. 19.

**Figure 19.**
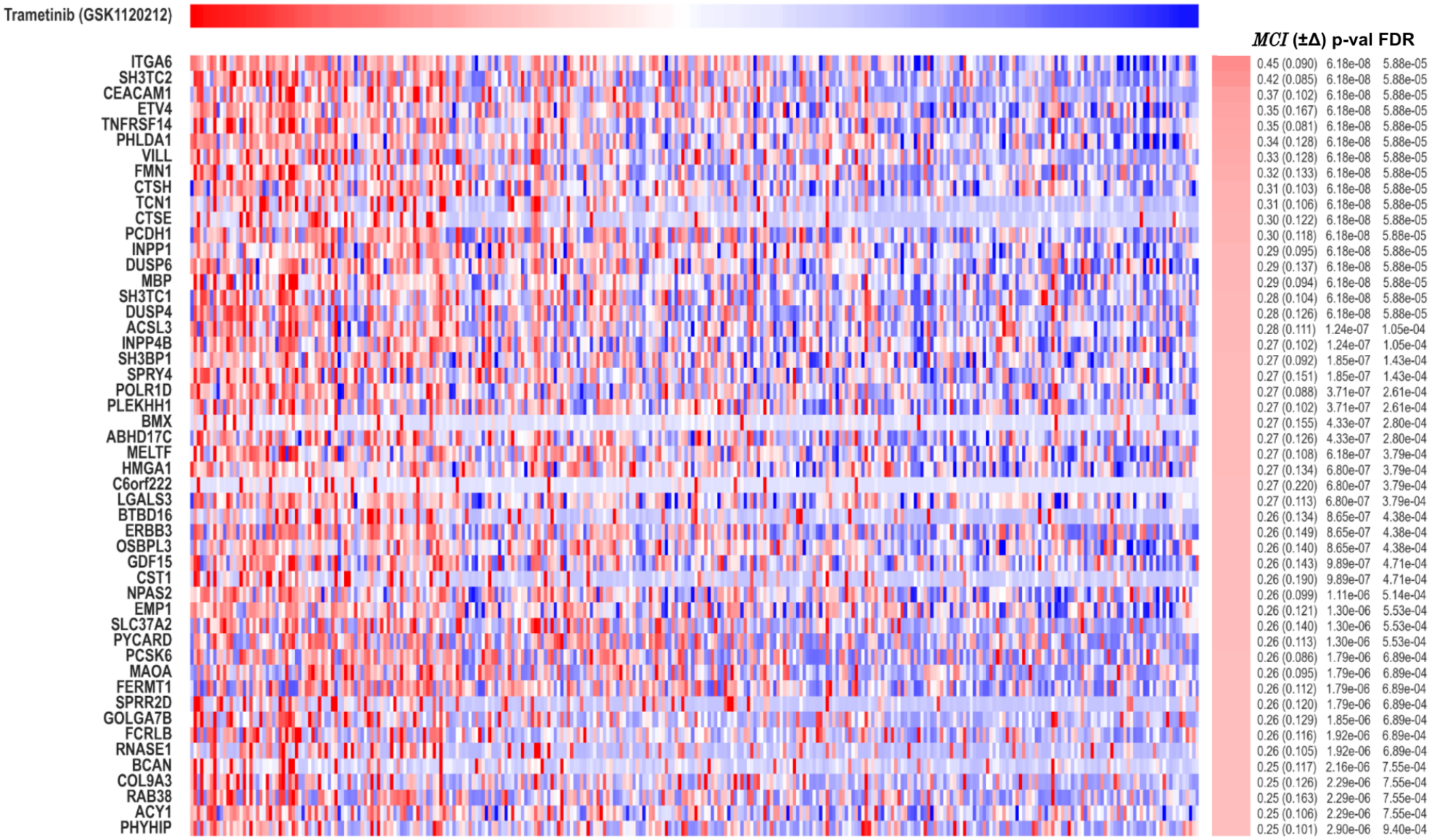
These are the top 50 gene profiles (mRNA) that associate with trametinib response using the CCLE mRNA and PRISM drug-screen datasets.

In order to generate a gene set representing the response to trametinib, we set an FDR threshold of 0.01 and find all the genes above that threshold (132 genes). This gene set can be used in combination with single-sample Gene Set Enrichment Analysis[71] (ssGSEA) to generate trametinib response signature scores in new cancer samples. We performed this procedure using the test set we defined before and we find that the ssGSEA scores using the *MCI* allows us to sort the CCLE test set of 158 cell lines according to their likelihood of response to Trametinib (Fig. 20). When we compare these scores vs. the actual response to Trametinib we can see that the signature scores are accurate and can be used as a biomarker. The *MCI* between the actual response and the signature profile is 0.57 a highly statistically significant match (p-value = 0.00001).

**Figure 20.**
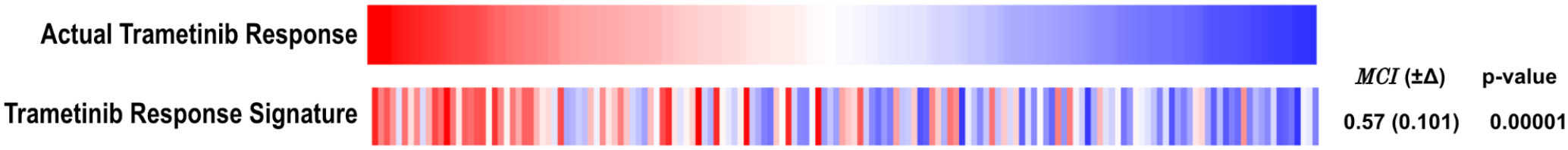
This figure shows the association between the actual response to trametinib in the 158 test set cell lines versus ssGSEA profiles of the *MCI*-generated signature (gene set).

As an additional application of the trametinib response signature, we generated ssGSEA scores for a group of 5 samples from melanoma patients treated with trametinib and dabrafenib and for which their transcriptional profiles were obtained before, during or after treatment [82] (Fig. 21). In all these cases the trametinib response signature scores decrease during or after treatment compared with its values for the same patient samples before treatment. This behavior indicates that the tumors are becoming less sensitive to the MAPK inhibition treatment as they overexpress immune PD-L1 positivity and immune activation signatures [82].

**Figure 21.**
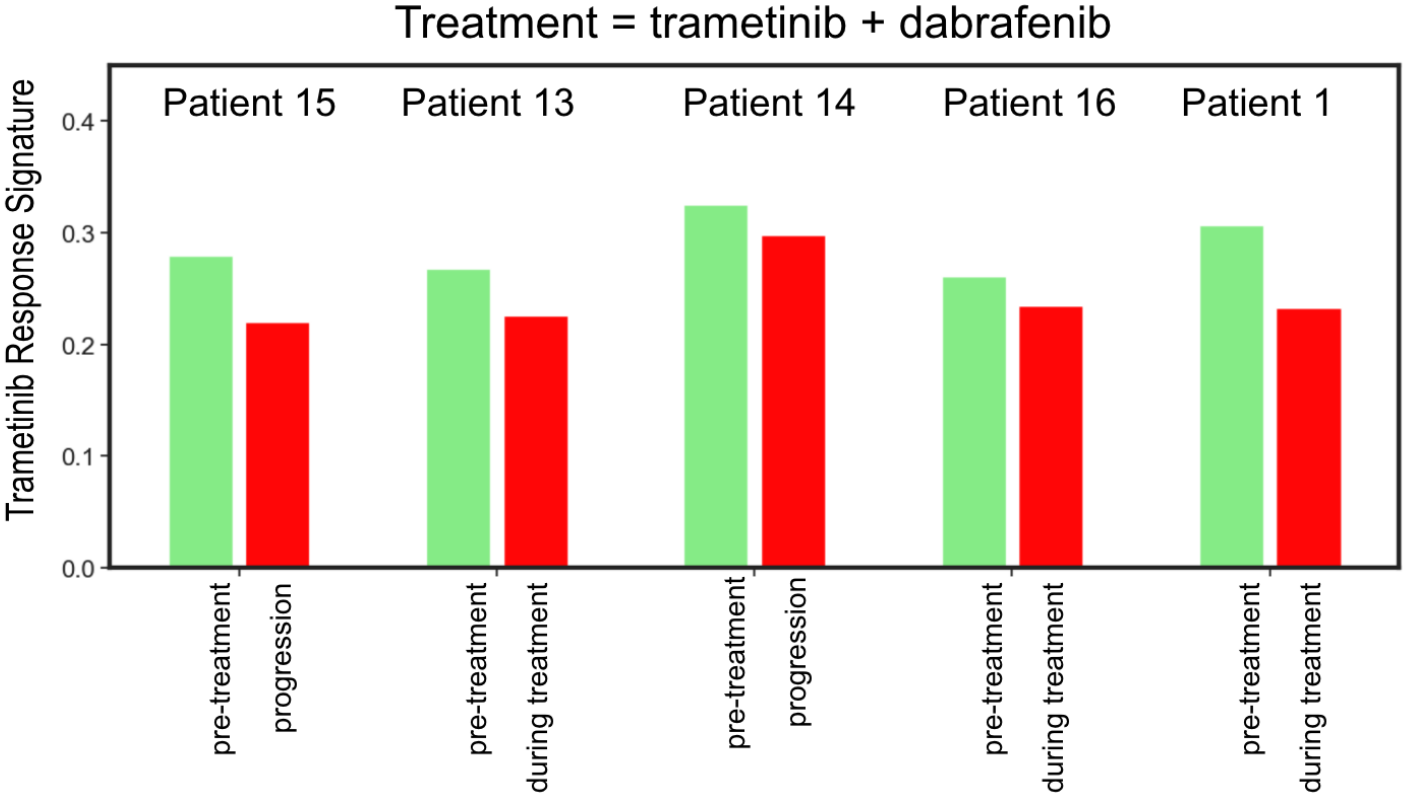
Decrease in trametinib response ssGSEA signature values when comparing profiles of melanoma patients before and during or after treatment with trametinib + dabrafenib.

#### Example 4: using the *MCI* to generate similarity matrices

In this example we show how the *MCI* can be used to generate similarity matrices for functional genomic datasets that in turn can be used for clustering or as input to network generating algorithms. The fact that the *MCI* can be computed efficiently allows us to process large datasets with *n* samples where each sample genomics profile, such as a vector of gene expression values, is compared against each other sample resulting in *n*^*2*^ computations of association. Fig. 22A below shows an *MCI*-based association matrix for the entire CCLE mRNA dataset that contains 1,304 cancer cell lines gene expression profiles for 16,184 genes. The computation of this matrix required 1,700,416 comparisons of 16,184-element long floating point vectors which took about one hour (3,606 secs) of computer time.^13^ Once generated the similarity matrix can be converted into a distance matrix, for example, by subtracting from one each *MCI* value so that the scale is inverted. In this way, similar sample profiles with *MCI* values near one become entries in the distance matrix with distances near zero. Entries with near zero *MCI*, will become entries with distances around one, and anticorrelated entries with *MCI* values near -1, will become entries with maximal distance near 2.

**Figure 22.**
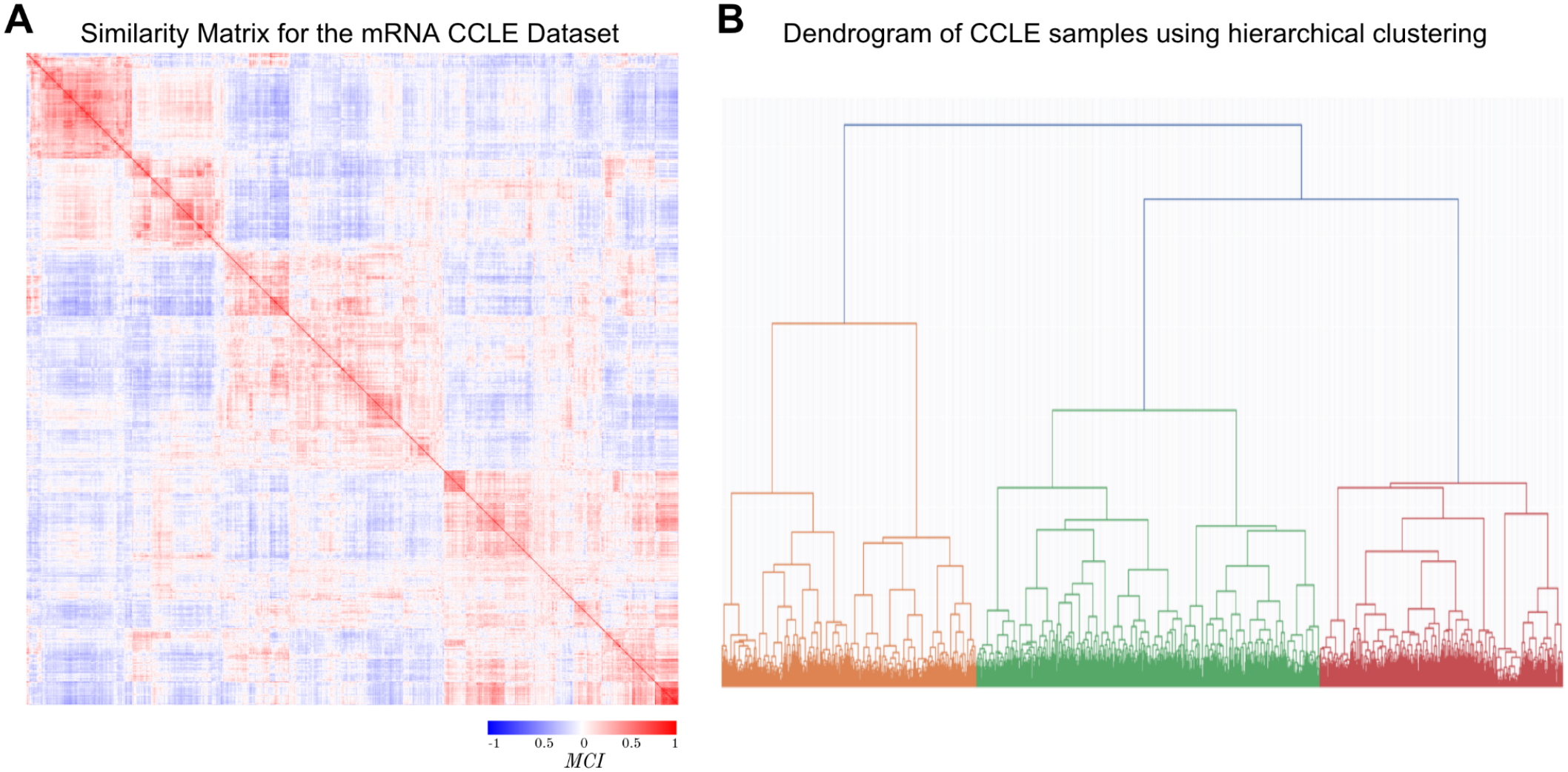
*MCI*-based association matrix for the CCLE mRNA dataset (A) and resulting dendrogram (B).

Once generated the distance matrix can be used to compute a clustering of the samples as shown in Fig. 22B, where a hierarchical clustering method, in this case *ward linkage*, was used to generate a dendrogram that organized the 1,304 samples according to their *MCI*-based distances. Analysis of this type using informational-theoretic association metrics in large genomic datasets is often limited because of long computation times. However, the *MCI* enables the possibility to perform information-based analysis of similarities in large genomic datasets due to its increased computational efficiency.

In the same way the *MCI* can be used to generate networks where the links represent the amount of association between nodes representing genomic variables [83]. This computational efficiency can enable the construction of genome-wide association networks that capture complex regulatory relationships, co-expression patterns, and functional modules across entire biological systems. These networks become the foundation for downstream analyses like pathway enrichment, disease gene prioritization, and drug target identification that would be impossible without efficient computation [84].

#### Example 5: Using the *MCI* to match binary and continuous variables

Here we show two examples of using the *MCI* and the Randomized Probability Integral Transform described above, where random Gaussian distributed noise is added to the binary variable in order to break the ties. The datasets are part of the Cancer Dependency Map (depmap.org). The first example shown in Fig. 23, shows how gene expression profiles can be matched against a binary phenotype in this case representing the mutation status of a well known driver oncogene: BRAF V600E. Among the top matches we can see the gene expression profile of BRAF itself and other MAPK genes and downstream relevant ones such as MAPK1, MITF, SOX10, ZEB2 and MAP2K1. The second example (Fig. 24) shows how binary features, in this case allele mutations, can be matched against a continuous phenotype, in this case the response to the BRAF-inhibitor dabrafenib. As one can see from the figure the top match is the mutation status of BRAF V600E. These two examples demonstrate how the MCI can be used effectively to match binary and continuous genomic variables.

**Figure 23.**
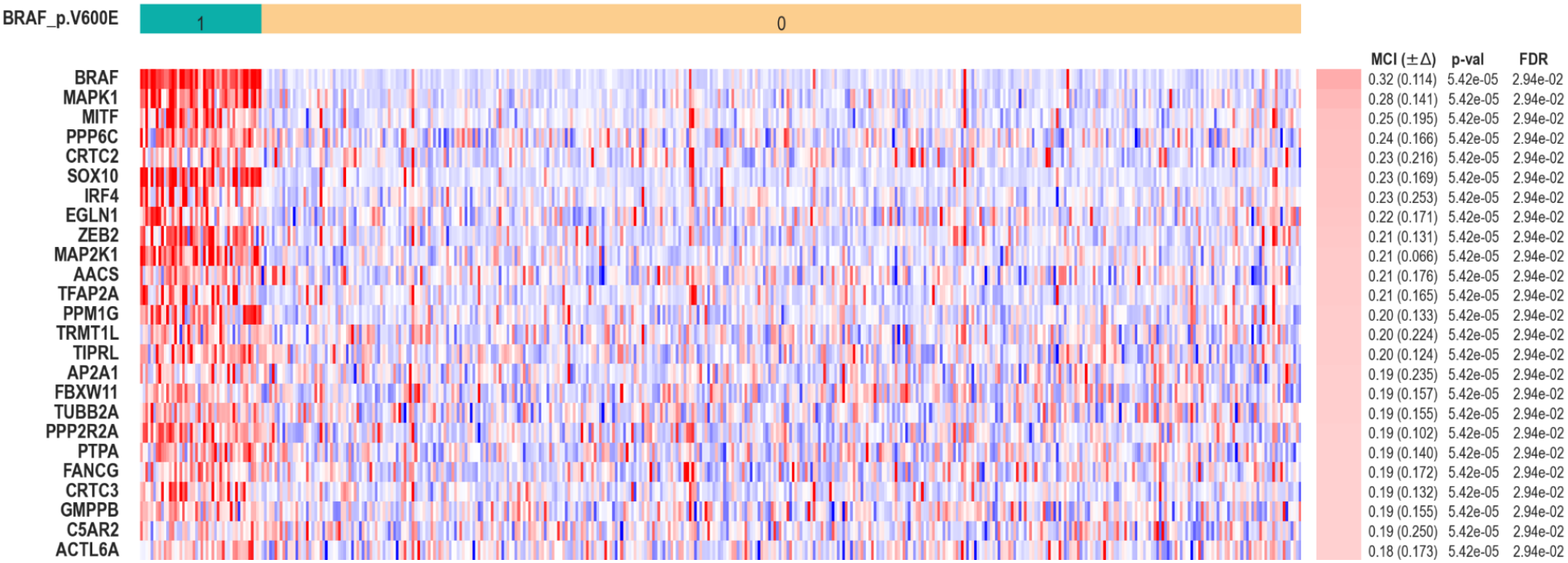
Matching continuous features (mRNA profiles) against a binary phenotype (mutation status).

**Figure 24.**
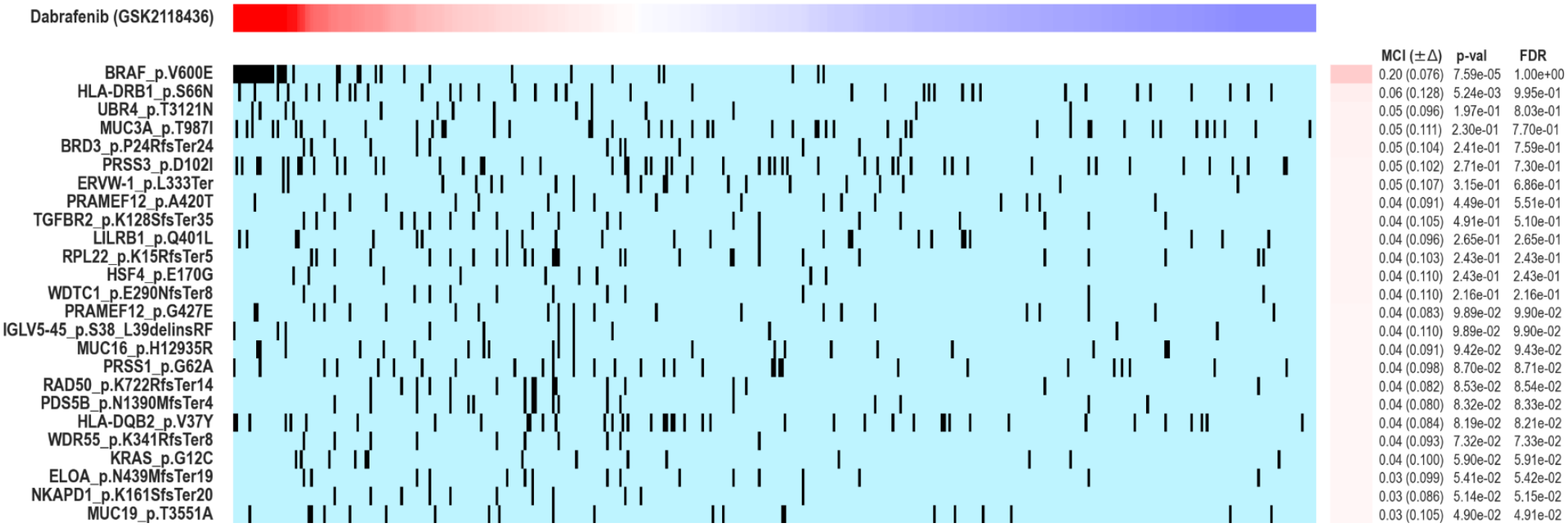
Matching binary features (allele mutations) against a continuous phenotype (dabrafenib response).

## Conclusions

We have introduced the *Multi-scale Copula Index* (*MCI*), a non-parametric rank-based association measure that works with empirical cumulative distribution functions, eliminating the need for density estimation or tuning parameters. As we have shown in several benchmarks, it can effectively capture complex non-linear dependencies, handles heavy-tailed distributions and cleanly separates dependence structure from marginal distributions. It is also effective to estimate association between genomic variables as we have illustrated in four examples, including estimating the degree of association of genes with protein expression, associating transcriptional and perturbational profiles of oncogenes, generating a drug response signature and generating similarity matrices or genome-wide association networks. The most important characteristics of the *MCI* can be summarized as follows:

1. **Empirical and non-parametric approach**: It does not make assumptions about the underlying data distributions, offering flexibility for analyzing diverse data structures. Additionally, it doesn’t require density estimation or internal parameter tuning as it works directly with empirical cumulative distribution functions rather than requiring kernel density estimation.
2. **Sample efficiency and estimation stability**: Because it is based on empirical cumulative distributions rather than densities, it typically requires fewer samples to achieve stable, reliable, and consistent estimates compared to mutual information, which relies on density estimation, especially with limited data. As a consequence of being rank-based and using the copula of cumulative distributions, it is less sensitive to outliers and extreme values.
3. **Computational efficiency**: The *MCI* rank-based calculations can be computed very efficiently, resulting in speed improvements of over 30 times compared to kernel-based mutual information estimation, making it comparable to the execution times of the Spearman correlation coefficient.
4. **Built-in directionality**: It provides a measure of directionality as it is intrinsically signed and able to detect and quantify both positive and negative associations between variables.
5. **Symmetry under ordering transformations**: It has desirable sign-reversal symmetries under reflection and complementation that make it suitable for many practical applications.
6. **Multi-scale sensitivity**: The *MCI* can detect associations in data at different scales.
7. **Non-linear dependencies**: It can effectively quantify complex non-linear dependencies that traditional correlation coefficients might miss or underestimate.
8. **Better handling of heavy-tailed distributions**: For heavy-tailed distributions where differential entropy can be difficult to estimate, the *MCI* provides stable estimation.
9. **Decoupling of marginal distribution effects**: The *MCI* separates dependence structure from marginal distributions, allowing cleaner analysis of multivariate relationships.
10. **Functional Cancer Genomics applications performance**: As we show below, the *MCI* performs well in practical functional genomic applications, including differential gene expression and matching genomic features versus functional readouts, such as drug sensitivities.

The *MCI* is an extension and improvement over other information-theoretic measures, including the Cumulative Copula Entropy (*CCE*) [14,15], the Information Coefficient (*IC*) [18,19], the Cumulative Entropy (*CE*) [41,42], and the Maximal Information Coefficient (*MIC*) [11]. The *MCI* incorporates several desirable properties and improvements over existing measures such as those while also possessing unique characteristics that make it particularly useful for addressing many challenges in functional genomics data analysis. This is especially relevant when genomic variables exhibit a distinct pattern of remaining uncorrelated across the majority of data points, while showing strong associations within specific subsets that represent “active” biological contexts as can be seen in the examples of Fig. 17. This prevalent phenomenon in functional genomics datasets presents an ideal opportunity to apply the *MCI* as a metric for quantifying these context-dependent associations. Its advantages render the *MCI* a valuable measure of dependency, particularly for applications involving continuous variables with complex statistical structures, such as non-trivial correlations, non-Gaussian or non-linear behaviors, limited sample sizes, or high-dimensional data, all common scenarios in functional cancer genomics data analysis. Additionally, the *MCI* is applicable to different variable types in other contexts beyond direct association and feature selection, including dimensionality reduction, matrix factorization, network reconstruction, pattern recognition, and predictive modeling.

## Acknowledgments

The work presented here was supported in part by NIH grants: R01CA154480, R01CA121941, R01DE026644, R01NS122835, R01CA257505, R01CA247551, R01DE026870, U24CA220341, U24CA248457, R01CA226803, R01CA292590, U01CA278470, U01CA268052, U01CA217885, P30CA023100, R01CA109467, R01DE035393, P01CA281819, U01CA176058, and a State of California’s *Initiative to Advance Precision Medicine* (CIAPM) award (OPR18112).

## Software

The Python and Cython programs that implement the computation of the *MCI*, and a Jupyter notebook with examples, are available at https://github.com/pablotamayo/MCI.

## Appendix

Effects of the Complementation Transformations on the MCI

We will analyze here in more detail the effect of the complementation transformation *U →* 1-*U*, for different components of the *MCI* entropy as a function of the different cumulative copulas and show that the MCI obeys the corresponding symmetries (eq. 24). Those symmetries require that under complementation of one of the input variables, the *MCI* will preserve its magnitude but change sign. As we mentioned in the main text the effect of these transformations is to reverse the ordering of the data along one of the cumulative copulas’ axes which results in pair exchanges of the values of the different *SCCE* in equation A1 below (eq. 28). We will focus on only one lattice but the argument can be applied to all the scales involved in the *MCI* computation. The *SCCE* (at one scale) is defined as,

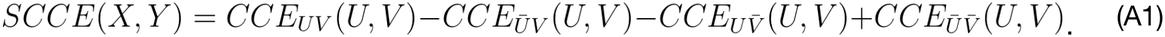

As the *CCEs*, and the entropulas in turn, are functions of the cumulative copulas we will study the cumulative copulas themselves (eq. 24),

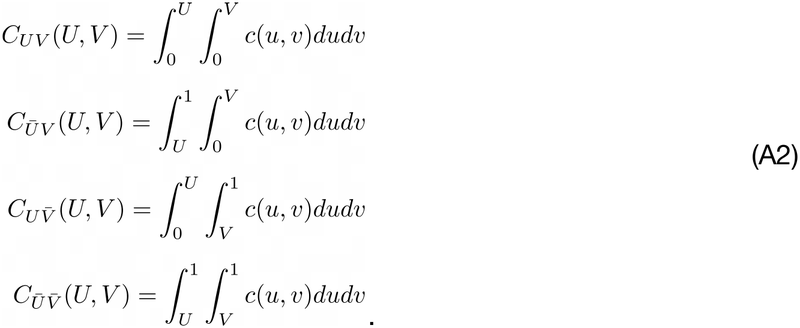

The second copula can be expanded and rearranged so its relation to the first copula is made apparent,

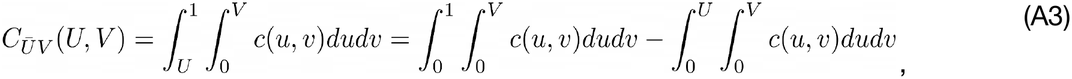

where the first term is only a function of *V* that will define as *ϕ(V)* and the last term is the first copula,

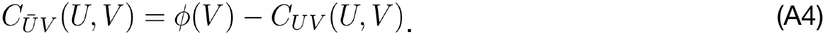

Analyzing the third copula in a similar way we obtain,

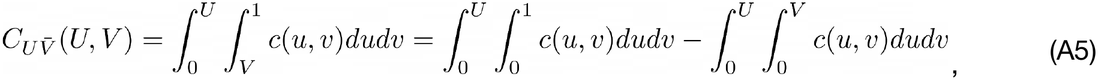

where the first term is only a function of *U* that we define as *ξ(U)* and the second term is the first copula.

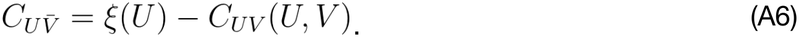

The last copula corresponds to,

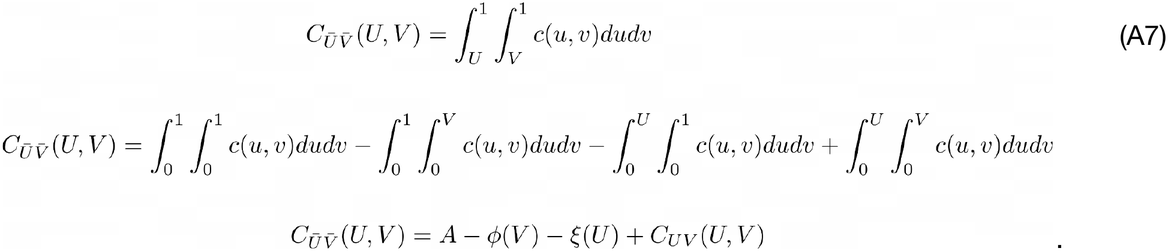

Where we used *ϕ(V)* and *ξ(U)* as before and introduced the constant *A*, for the first integral that corresponds to the area under the copula. In summary, the 4 copulas as functions of *C*_*UV*_*(U, V), ϕ(V), ξ(U)* and *A*, are,

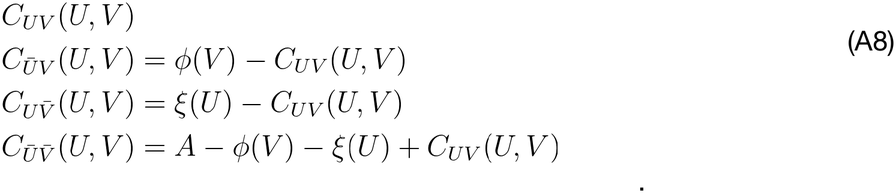

Now let’s analyze the effect of the complementation *U →* 1-*U* on these copulas. Analyzing the first copula, we can see that the effect of complementation on *U* amounts to changing the sign of *C*_*UV*_*(U,V)* but also introducing an additional *ϕ(V)* factor,

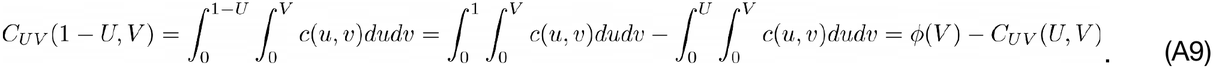

This explains why the *CCE* is not symmetric under complementation. A similar computation shows the effect of complementation on the other copulas,

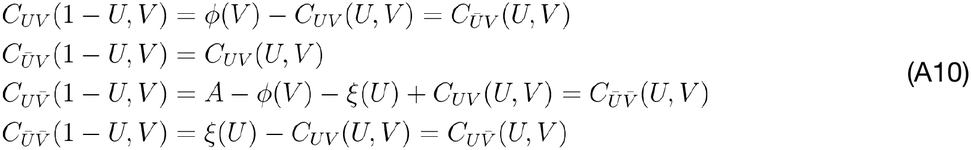

where we can see that the complementation, as expected, exchanges the values of the first two and the last two copulas. Then when we compute the *SCCE*, the result of this exchange will amount to changing the sign of each term in eq. A1 and therefore an overall change of sign of the *SCCE*, and consequently the same change in sign for the *MSCCE*, and the *RMSCCE* (*MCI*) but leaving its magnitude unchanged as required. Similar derivations apply to the *V →* 1-*V* complementation and the *V →* -*V* and *U →* -*U* reflections.

## Footnotes

1 The CRISPR and mRNA datasets used in these examples correspond to release 25Q3 of the DepMap. In this analysis the CRISPR profiles have been reflected, i.e. taken with negative signs so that higher sensitivity to knockdown corresponds to higher values of the profile. The resulting values were also standardized and clipped at 7 standard deviations above/below the mean. The mRNA profiles (x-axis) are also standardized in the same way.

2 In order to accurately estimate the extremely low p-values for the associations shown in Figs. 3 and 4 we made use of the Wang-Landau algorithm [85,86] to efficiently compute the tails of the measures’ empirical null distribution and enable the estimation of small p-values using adaptive importance sampling.

3 Our implementation of the *IC* makes use of the bandwidth (*bcv*) and kernel density estimation (*kde2d*) functions from R’s package *MASS* and adjusting the bandwidth estimate to take into account the correlation between vectors where the kernel density bandwidth is bcv_bandwidth_ x (1 - |¾ *ϱ* (x,y)|), where *ϱ*; is the pearson correlation.

4 Sklar chose the latin word because a “copula” functions logically as a “linkage”, it physically couples or joins a multivariate joint cumulative distribution function to its individual one-dimensional marginal distribution functions. Two-dimensional copulas are known in some other areas of mathematics under the name *permutons* and *doubly-stochastic* measures.

5 In this equation the 1(…) are indicator functions that return the value 1 if the expression in parenthesis is true and 0 if the expression in parentheses is false

6 The heuristic of using odd exponential power terms is based on the fact that an analytical mapping between the normalized MCI measure and Spearman’s will have to be an odd function, strictly monotonic, smooth, and pinned at three exact points (-1, 0, 1). The power–exponential structure, rather than standard polynomial, is motivated by Linfoot’s informational coefficient [19] and the Gaussian-copula relationship between latent correlation and Spearman’s rho [26].

7 It appears to be faster than the computation of the Pearson correlation coefficient using the *scipy*.*stats*.*pearsonr* function for vectors of size less than about 800 data points.

8 Anscombe’s quartet is a famous dataset in statistics and data science [87] that consists of four datasets that have nearly identical descriptive statistics (mean, variance, correlation), yet appear very different when graphed.

9 This is achieved by generating *n* samples from a bivariate Gaussian distribution with pre-specified means and cross-correlation using the *random*.*multivariate_normal* function from the *numpy* python package.

10 The timings were obtained from a *MacBook Pro* computer (M1 3.2 GHz CPU with 64 GB of memory).

11 Reverse Phase Protein Array (RRPA) are an antibody-based high throughput proteomic approach that enables the simultaneous quantification of large sample number of proteins [88] is shown in Fig. 14 where the *MCI* decays rapidly as one moves down the list of genes and at the bottom of the list there are genes that display a strong negative association.

12 The p-values were computed by an empirical permutation test that generated random scores where 10^6^ random permutations of the mRNA profiles were compared against the observed CRISPR profile. The p-values were computed from the ratio of the number of random scores equal or higher than the observed score divided by the number of random permutations.

13 The computer execution timings were obtained in a MacBook Pro computer (M1 3.2 GHz CPU with 64 GB of main memory).

## References

1. Mohr AE, Ortega-Santos CP, Whisner CM, Klein-Seetharaman J, Jasbi P. Navigating Challenges and Opportunities in Multi-Omics Integration for Personalized Healthcare. Biomedicines. 2024;12. doi:10.3390/biomedicines12071496

2. Li R, Li L, Xu Y, Yang J. Machine learning meets omics: applications and perspectives. Brief Bioinform. 2022;23. doi:10.1093/bib/bbab460

3. Wolde T, Bhardwaj V, Pandey V. Current Bioinformatics Tools in Precision Oncology. MedComm (2020). 2025;6: e70243.

4. Cominetti O, Moreno SO, Agarwal S. Advances in methods and tools for multi-omics data analysis. Frontiers Media SA; 2023.

5. Mukherjee A, Abraham S, Singh A, Balaji S, Mukunthan KS. From Data to Cure: A Comprehensive Exploration of Multi-omics Data Analysis for Targeted Therapies. Mol Biotechnol. 2025;67: 1269–1289.

6. Hahn WC, Bader JS, Braun TP, Califano A, Clemons PA, Druker BJ, et al. An expanded universe of cancer targets. Cell. 2021;184: 1142–1155.

7. Boehm JS, Hahn WC. Towards systematic functional characterization of cancer genomes. Nat Rev Genet. 2011;12: 487–498.

8. Tamayo P, Steinhardt G, Liberzon A, Mesirov JP. The limitations of simple gene set enrichment analysis assuming gene independence. Stat Methods Med Res. 2016;25: 472–487.

9. Székely GJ, Rizzo ML, Bakirov NK. Measuring and testing dependence by correlation of distances. Ann Stat. 2007;35: 2769–2794.

10. Wang T, Dai X, Liu Y. Learning with Hilbert–Schmidt independence criterion: A review and new perspectives. Knowl Based Syst. 2021;234: 107567.

11. Reshef DN, Reshef YA, Finucane HK, Grossman SR, McVean G, Turnbaugh PJ, et al. Detecting novel associations in large data sets. Science. 2011;334: 1518–1524.

12. Moon YI, Rajagopalan B, Lall U. Estimation of mutual information using kernel density estimators. Phys Rev E Stat Phys Plasmas Fluids Relat Interdiscip Topics. 1995;52: 2318–2321.

13. Kraskov A, Stögbauer H, Grassberger P. Estimating mutual information. Phys Rev E Stat Nonlin Soft Matter Phys. 2004;69: 066138.

14. Sunoj SM, Unnikrishnan Nair N. Survival Copula Entropy and Dependence in Bivariate Distributions. REVSTAT. 2025;23: 101–115.

15. Arshad M, Zachariah SG, Pathak AK. Multivariate information measures: A copula-based approach. arXiv [stat.ME]. 2024. Available: http://arxiv.org/abs/2408.02028

16. Chang HR, Jung E, Cho S, Jeon Y-J, Kim Y. Targeting Non-Oncogene Addiction for Cancer Therapy. Biomolecules. 2021;11. doi:10.3390/biom11020129

17. Weinstein IB, Joe A. Oncogene addiction. Cancer Res. 2008;68: 3077–80; discussion 3080.

18. Joe H. Relative entropy measures of multivariate dependence. J Am Stat Assoc. 1989;84: 157–164.

19. Linfoot EH. An informational measure of correlation. Information and Control. 1957;1: 85–89.

20. Ma JY, Ting S, Tam B, Pham T, Reich M, Mesirov J, et al. Deciphering the Functional Roles of Individual Cancer Alleles Across Comprehensive Cancer Genomic Studies. bioRxiv. 2023. doi:10.1101/2023.11.14.567106

21. Kim JW, Botvinnik OB, Abudayyeh O, Birger C, Rosenbluh J, Shrestha Y, et al. Characterizing genomic alterations in cancer by complementary functional associations. Nat Biotechnol. 2016;34: 539–546.

22. Abazeed ME, Adams DJ, Hurov KE, Tamayo P, Creighton CJ, Sonkin D, et al. Integrative radiogenomic profiling of squamous cell lung cancer. Cancer Res. 2013;73: 6289–6298.

23. Cowley GS, Weir BA, Vazquez F, Tamayo P, Scott JA, Rusin S, et al. Parallel genome-scale loss of function screens in 216 cancer cell lines for the identification of context-specific genetic dependencies. Sci Data. 2014;1: 140035.

24. Liberzon A, Birger C, Thorvaldsdóttir H, Ghandi M, Mesirov JP, Tamayo P. The Molecular Signatures Database (MSigDB) hallmark gene set collection. Cell Syst. 2015;1: 417–425.

25. Sarto Basso R, Hochbaum DS, Vandin F. Efficient algorithms to discover alterations with complementary functional association in cancer. PLoS Comput Biol. 2019;15: e1006802.

26. Nelsen RB. An Introduction to Copulas. Springer Science & Business Media; 2007.

27. Joe H. Dependence Modeling with Copulas. CRC Press; 2014.

28. Durante F, Sempi C. Principles of Copula Theory. CRC Press; 2015.

29. Cover TM, Thomas JA. Elements of Information Theory. John Wiley & Sons; 2006.

30. Roman S. Introduction to Coding and Information Theory. Springer Science & Business Media; 1996.

31. Polyanskiy Y, Wu Y. Information Theory: From Coding to Learning. Cambridge University Press; 2025.

32. Banerjee S, Yoon H, Ting S, Tang C-M, Yebra M, Wenzel AT, et al. KITlow Cells Mediate Imatinib Resistance in Gastrointestinal Stromal Tumor. Mol Cancer Ther. 2021;20: 2035–2048.

33. Ren S, Gaykalova DA, Guo T, Favorov AV, Fertig EJ, Tamayo P, et al. HPV E2, E4, E5 drive alternative carcinogenic pathways in HPV positive cancers. Oncogene. 2020;39: 6327–6339.

34. Wu X, Yeerna H, Goto Y, Ando T, Wu VH, Zhang X, et al. Metformin Inhibits Progression of Head and Neck Squamous Cell Carcinoma by Acting Directly on Carcinoma-Initiating Cells. Cancer Res. 2019;79: 4360–4370.

35. Reilly BM, Tanaka TN, Diep D, Yeerna H, Tamayo P, Zhang K, et al. DNA Methylation Identifies Genetically and Prognostically Distinct Subtypes of MDS. Blood. 2018. pp. 106–106. doi:10.1182/blood-2018-99-111535

36. Deichaite I, Sears TJ, Sutton L, Rebibo D, Morgan K, Nelson T, et al. Differential regulation of TNFα and IL-6 expression contributes to immune evasion in prostate cancer. J Transl Med. 2022;20: 527.

37. Huang FW, Mosquera JM, Garofalo A, Oh C, Baco M, Amin-Mansour A, et al. Exome Sequencing of African-American Prostate Cancer Reveals Loss-of-Function Mutations. Cancer Discov. 2017;7: 973–983.

38. Hanaford AR, Archer TC, Price A, Kahlert UD, Maciaczyk J, Nikkhah G, et al. DiSCoVERing Innovative Therapies for Rare Tumors: Combining Genetically Accurate Disease Models with In Silico Analysis to Identify Novel Therapeutic Targets. Clin Cancer Res. 2016;22: 3903–3914.

39. Wood KC, Konieczkowski DJ, Johannessen CM, Boehm JS, Tamayo P, Botvinnik OB, et al. MicroSCALE screening reveals genetic modifiers of therapeutic response in melanoma. Sci Signal. 2012;5: rs4.

40. Konieczkowski DJ, Johannessen CM, Abudayyeh O, Kim JW, Cooper ZA, Piris A, et al. A melanoma cell state distinction influences sensitivity to MAPK pathway inhibitors. Cancer Discov. 2014;4: 816–827.

41. Di Crescenzo A, Longobardi M. On cumulative entropies. J Stat Plan Inference. 2009;139: 4072–4087.

42. Rao M, Chen Y, Vemuri BC, Wang F. Cumulative residual entropy: A new measure of information. IEEE Trans Inf Theory. 2004;50: 1220–1228.

43. Wang F, Vemuri BC, Rao M, Chen Y. A new & robust information theoretic measure and its application to image alignment. Inf Process Med Imaging. 2003;18: 388–400.

44. Ma J, Sun Z. Mutual information is copula entropy. Tsinghua Sci Technol. 2011;16: 51–54.

45. Calsaverini RS, Vicente R. An information theoretic approach to statistical dependence: copula information. arXiv [q-fin.ST]. 2009. Available: http://arxiv.org/abs/0911.4207

46. Kumar N, Dixit A, Vijay V. Entropy measures and their applications: A comprehensive review. arXiv [math.PR]. 2025. Available: http://arxiv.org/abs/2503.15660

47. Loeve M. Probability Theory: Third Edition. Courier Dover Publications; 2017.

48. Klein I, Doll M. (Generalized) Maximum Cumulative Direct, Residual, and Paired Φ Entropy Approach. Entropy (Basel). 2020;22. doi:10.3390/e22010091

49. Brower RC, Giles R, Moriarty KJM, Tamayo P. Combining renormalization group and multigrid methods. J Comput Phys. 1989;80: 472–479.

50. Brower RC, Tamayo P, York B. A parallel multigrid algorithm for percolation clusters. J Stat Phys. 1991;63: 73–88.

51. Sunoj SM, Unnikrishnan Nair N. Survival Copula Entropy and Dependence in Bivariate Distributions. REVSTAT. 2025;23: 101–115.

52. Dunn PK, Smyth GK. Randomized Quantile Residuals. J Comput Graph Stat. 1996;5: 236–244.

53. Correa CD, Lindstrom P. The mutual information diagram for uncertainty visualization. Int J Uncertain Quantif. 2013;3: 187–201.

54. Taylor KE. Summarizing multiple aspects of model performance in a single diagram. J Geophys Res. 2001;106: 7183–7192.

55. Rosas FE, Mediano PAM, Gastpar M, Jensen HJ. Quantifying high-order interdependencies via multivariate extensions of the mutual information. Phys Rev E. 2019;100: 032305.

56. Lizier JT, Bertschinger N, Jost J, Wibral M. Information Decomposition of Target Effects from Multi-Source Interactions: Perspectives on Previous, Current and Future Work. Entropy (Basel). 2018;20. doi:10.3390/e20040307

57. Huang JK, Carlin DE, Yu MK, Zhang W, Kreisberg JF, Tamayo P, et al. Systematic Evaluation of Molecular Networks for Discovery of Disease Genes. Cell Syst. 2018;6: 484–495.e5.

58. Carter H, Hofree M, Ideker T. Genotype to phenotype via network analysis. Curr Opin Genet Dev. 2013;23: 611–621.

59. Barretina J, Caponigro G, Stransky N, Venkatesan K, Margolin AA, Kim S, et al. The Cancer Cell Line Encyclopedia enables predictive modelling of anticancer drug sensitivity. Nature. 2012;483: 603–607.

60. Cheung HW, Cowley GS, Weir BA, Boehm JS, Rusin S, Scott JA, et al. Systematic investigation of genetic vulnerabilities across cancer cell lines reveals lineage-specific dependencies in ovarian cancer. Proc Natl Acad Sci U S A. 2011;108: 12372–12377.

61. Speyer G, Mahendra D, Tran HJ, Kiefer J, Schreiber SL, Clemons PA, et al. DIFFERENTIAL PATHWAY DEPENDENCY DISCOVERY ASSOCIATED WITH DRUG RESPONSE ACROSS CANCER CELL LINES. Pac Symp Biocomput. 2017;22: 497–508.

62. Seashore-Ludlow B, Rees MG, Cheah JH, Cokol M, Price EV, Coletti ME, et al. Harnessing Connectivity in a Large-Scale Small-Molecule Sensitivity Dataset. Cancer Discov. 2015;5: 1210–1223.

63. Rees MG, Seashore-Ludlow B, Clemons PA. Computational Analyses Connect Small-Molecule Sensitivity to Cellular Features Using Large Panels of Cancer Cell Lines. Methods Mol Biol. 2019;1888: 233–254.

64. Cokelaer T, Chen E, Iorio F, Menden MP, Lightfoot H, Saez-Rodriguez J, et al. GDSCTools for Mining Pharmacogenomic Interactions in Cancer. 2017. doi:10.1101/166223

65. Iorio F, Knijnenburg TA, Vis DJ, Bignell GR, Menden MP, Schubert M, et al. A Landscape of Pharmacogenomic Interactions in Cancer. Cell. 2016;166: 740–754.

66. Yang W, Lightfoot H, Bignell G, Behan F, Cokelear T, Haber D, et al. Genomics of Drug Sensitivity in Cancer (GDSC): A resource for biomarker discovery in cancer cells. Eur J Cancer. 2016;69: S82.

67. Subramanian A, Narayan R, Corsello SM, Peck DD, Natoli TE, Lu X, et al. A Next Generation Connectivity Map: L1000 Platform and the First 1,000,000 Profiles. Cell. 2017;171: 1437–1452.e17.

68. Boehm JS, Golub TR. An ecosystem of cancer cell line factories to support a cancer dependency map. Nat Rev Genet. 2015;16: 373–374.

69. Krogan NJ, Lippman S, Agard DA, Ashworth A, Ideker T. The Cancer Cell Map Initiative: Defining the Hallmark Networks of Cancer. Mol Cell. 2015;58: 690–698.

70. Network TCTDAD, The Cancer Target Discovery and Development Network. Transforming Big Data into Cancer-Relevant Insight: An Initial, Multi-Tier Approach to Assess Reproducibility and Relevance. Molecular Cancer Research. 2016. pp. 675–682. doi:10.1158/1541-7786.mcr-16-0090

71. Scholl C, Fröhling S, Dunn IF, Schinzel AC, Barbie DA, Kim SY, et al. Synthetic lethal interaction between oncogenic KRAS dependency and STK33 suppression in human cancer cells. Cell. 2009;137: 821–834.

72. Barbie DA, Tamayo P, Boehm JS, Kim SY, Moody SE, Dunn IF, et al. Systematic RNA interference reveals that oncogenic KRAS-driven cancers require TBK1. Nature. 2009;462: 108–112.

73. Papageorgiou A, Rapley J, Mesirov JP, Tamayo P, Avruch J. A genome-wide siRNA screen in mammalian cells for regulators of S6 phosphorylation. PLoS One. 2015;10: e0116096.

74. Paolella BR, Gibson WJ, Urbanski LM, Alberta JA, Zack TI, Bandopadhayay P, et al. Copy-number and gene dependency analysis reveals partial copy loss of wild-type SF3B1 as a novel cancer vulnerability. Elife. 2017;6. doi:10.7554/eLife.23268

75. Viswanathan SR, Nogueira MF, Buss CG, Krill-Burger JM, Wawer MJ, Malolepsza E, et al. Genome-scale analysis identifies paralog lethality as a vulnerability of chromosome 1p loss in cancer. Nat Genet. 2018;50: 937–943.

76. Kryukov GV, Wilson FH, Ruth JR, Paulk J, Tsherniak A, Marlow SE, et al. MTAP deletion confers enhanced dependency on the PRMT5 arginine methyltransferase in cancer cells. Science. 2016;351: 1214–1218.

77. Barbie TU, Alexe G, Aref AR, Li S, Zhu Z, Zhang X, et al. Targeting an IKBKE cytokine network impairs triple-negative breast cancer growth. J Clin Invest. 2014;124: 5411–5423.

78. Kim E, Ilic N, Shrestha Y, Zou L, Kamburov A, Zhu C, et al. Systematic Functional Interrogation of Rare Cancer Variants Identifies Oncogenic Alleles. Cancer Discov. 2016;6: 714–726.

79. Good P. Permutation Tests: A Practical Guide to Resampling Methods for Testing Hypotheses. Springer Science & Business Media; 2013.

80. Azzalini A. The skew-normal distribution and related multivariate families. Scand Stat Theory Appl. 2005;32: 159–188.

81. Yu C, Mannan AM, Yvone GM, Ross KN, Zhang Y-L, Marton MA, et al. High-throughput identification of genotype-specific cancer vulnerabilities in mixtures of barcoded tumor cell lines. Nat Biotechnol. 2016;34: 419–423.

82. Kakavand H, Rawson RV, Pupo GM, Yang JYH, Menzies AM, Carlino MS, et al. PD-L1 Expression and Immune Escape in Melanoma Resistance to MAPK Inhibitors. Clin Cancer Res. 2017;23: 6054–6061.

83. Yan J, Risacher SL, Shen L, Saykin AJ. Network approaches to systems biology analysis of complex disease: integrative methods for multi-omics data. Brief Bioinform. 2018;19: 1370–1381.

84. Guala D, Ogris C, Müller N, Sonnhammer ELL. Genome-wide functional association networks: background, data & state-of-the-art resources. Brief Bioinform. 2020;21: 1224–1237.

85. Wang F, Landau DP. Efficient, multiple-range random walk algorithm to calculate the density of states. Phys Rev Lett. 2001;86: 2050–2053.

86. Landau DP, Binder K. A Guide to Monte Carlo Simulations in Statistical Physics. Cambridge University Press; 2014.

87. Tufte E. The visual display of quantitative information (1983). In: Schneider B, Ernst C, Wöpking J, editors. Diagrammatik-Reader. Berlin, Boston: De Gruyter; 2016. pp. 219–230.

88. Coarfa C, Grimm SL, Rajapakshe K, Perera D, Lu H-Y, Wang X, et al. Reverse-Phase Protein Array: Technology, Application, Data Processing, and Integration. J Biomol Tech. 2021;32: 15–29.

